# Photosensitivity of an aging brain

**DOI:** 10.1101/2025.09.21.677632

**Authors:** Stephen Thankachan, Dmitriy Atochin, Ksenia Kastanenka, Dmitry Gerashchenko

**Affiliations:** Department of Psychiatry, Harvard Medical School and Veterans Affairs Boston Healthcare System, West Roxbury, MA, 02132, USA; Department of Neurology, Massachusetts General Hospital, Harvard Medical School, Charlestown, MA, 02129, USA

## Abstract

Red light is considered less phototoxic than blue light and is widely used in both research and photobiomodulation therapy. The difference in the response to brain exposure to light between young and old mammals is currently unknown. We found that brain exposure to blue light caused local damage in the cerebral cortex in both young and old mice. Brain exposure to red light did not have any noticeable effect on young mice. However, it caused a marked reduction in electroencephalogram power, damaged fiber bundles throughout the brain, and brought about a coma-like state in old mice. The effect of red light on electroencephalogram power was dose-dependent and particularly strong in the theta range. When delivered at a lower intensity but over a longer period, red light produced a similar reduction in electroencephalogram power and brain damage as those seen in the mice treated with higher irradiation over a shorter period. These results indicate that the impact of light on electroencephalogram and brain tissue strongly depends not only on the light wavelength, duration and intensity of the exposure, but also on the age of the animal and type of tissue exposed to light.

## Introduction

Photostimulation of brain tissue is widely used in both research and therapies. Light of shorter wavelengths, such as blue or green, has been commonly used in optogenetic studies^1^. Recently, red-light optogenetic tools have been developed and are gaining increasingly popularity^2^. The reasons for using red light (620-750 nm) in neuroscience include a better penetration of light of a longer wavelength into deep brain regions, low phototoxicity to neurons, and no significant impact on gene expression in brain cells compared to blue light (450-495 nm) delivered at the same intensity^2–4^. Red light is also used in photobiomodulation and other forms of red light therapy to help skin, muscle tissue, and other parts of the body heal^5, 6^. The LED panels and helmets for performing red light therapy in a home setting are freely available on the market.

Optogenetic studies are typically performed in young laboratory animals. Although investigators also performed experiments on older animals, light was delivered locally via a fiber-optic cannula that irradiated only a few mm^3^ of brain tissue^7–10^. To the best of our knowledge, the effects of light irradiation on the entire brain, or even a considerably large brain area, have not been investigated in mammals. A recent study demonstrated that susceptibility to blue light stress was age-dependent in Drosophila^11^. Blue light of the same intensity and duration reduced survival and increased neurodegeneration to a greater extent in old flies compared to young flies^11^. These results suggest that there may also be differences in the brain response to light exposure between young and old mammals. Considering that light irradiation of large brain areas is common, including exposure to sunlight and photobiomodulation therapies, it is important to understand the effects of light irradiation on the mammalian brain at different ages.

In the present study, we determined the effects of light irradiation on brain damage as a function of age and estimated the functional consequences of that damage on sleep. To that end, we exposed a large area of the brain to light by placing an LED on the top of the mouse skull. These experiments were performed on animals of different ages using both blue- and red-light LEDs. We hypothesized that the effect of light exposure on the brain depends on the light wavelength and the age of the animal.

## Results

### Effect of blue vs red light on young mice

We first compared the effect of blue vs red light in young animals. Three-month-old C57BL/6 mice were implanted with an LED on the top of the skull as well as electroencephalogram (EEG) and electromyogram (EMG) electrodes (Fig. 1A). Either red or blue light was delivered in 10 ms pulses at a frequency of 10 Hz and with a power of about 15 mW (red light) or 10 mW (blue light) during non-rapid eye movement (NREM) sleep using a closed-loop stimulation system^12^. We assessed whether 8.5 hours of such light stimulation delivered during the period of 24 hours impaired sleep and caused brain damage.

**Fig. 1.**
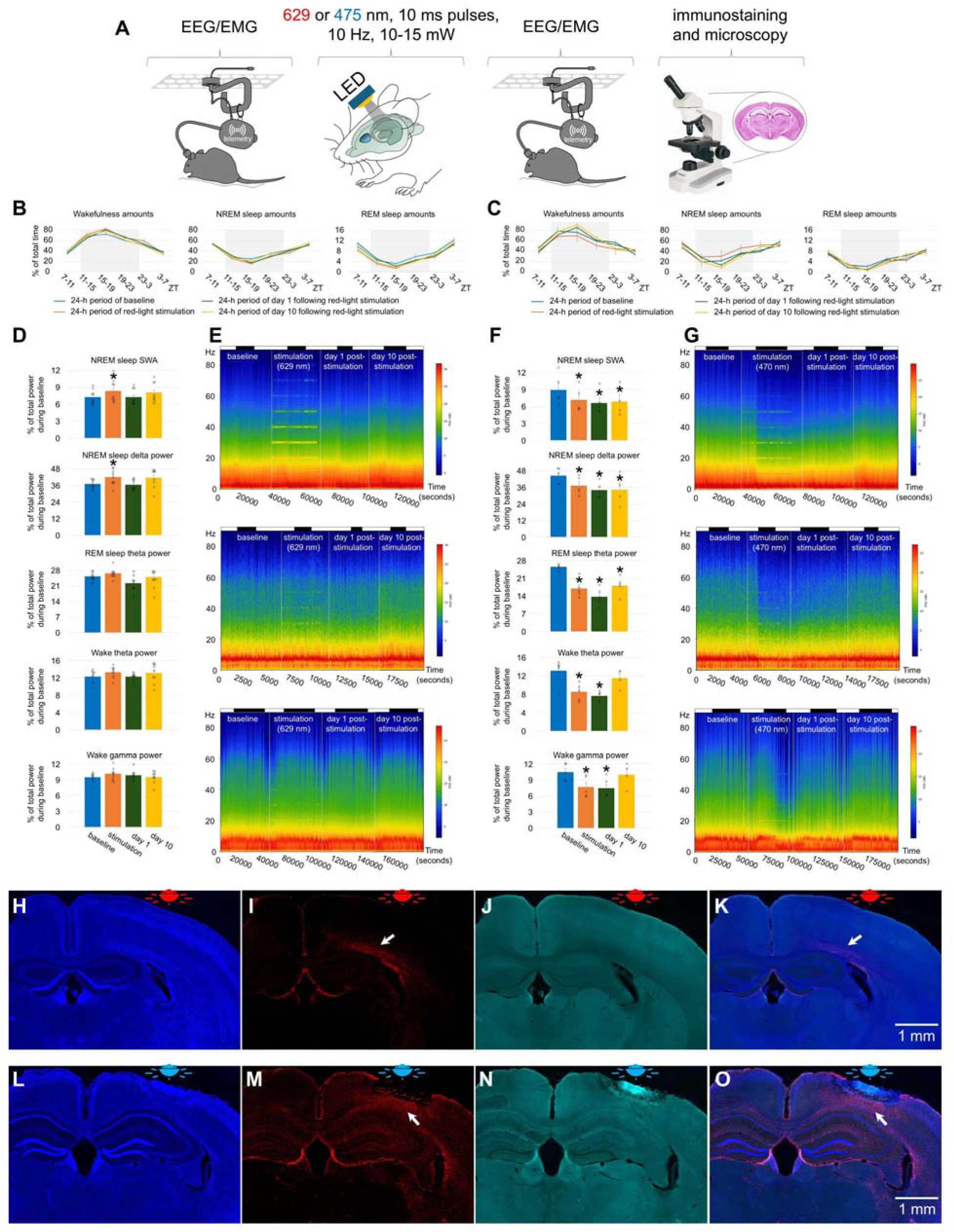
Effect of red and blue light on the brain of young mice. (**A**) Experimental designs. (**B**) Amounts of sleep and wakefulness in mice exposed to red light. (**C**) Amounts of sleep and wakefulness in mice exposed to blue light. (**D**) EEG power in SWA, delta, theta and gamma ranges in mice exposed to red light. (**E**) Representative heatmap of EEG power spectrum during NREM sleep (Top), REM sleep (Middle), and wakefulness (Bottom) in a mouse exposed to red light. (**F**) EEG power in SWA, delta, theta and gamma ranges in mice exposed to blue light. One-way repeated measures ANOVA followed by Tukey’s post hoc test, \**P* < 0.05 vs baseline. (**G**) Representative heatmap of EEG power spectrum during NREM sleep (Top), REM sleep (Middle), and wakefulness (Bottom) in a mouse exposed to blue light. (**H**) DAPI staining, (**I**) GFAP staining, (**J**) autofluorescence under a green filter, and (**K**) merged image of H-J in a mouse exposed to red light. (**L**) DAPI staining, (**M**) GFAP staining, (**N**) autofluorescence under a green filter, and (**O**) merged image of L-N in a mouse exposed to blue light. Error bars indicate the standard error of the mean. A bar on the top of figures A and G indicates light on and light off periods.

Red light stimulation did not result in a significant change in the amount of wakefulness and sleep (Fig. 1B). Estimation of EEG spectral density by multitaper analysis also did not reveal any significant changes except a transient increase within low frequency range during stimulation (Fig. 1D, Extended Data Fig. 1A and 1B). Notably, a peak at the fundamental frequency of 10 Hz and harmonics at 10 Hz intervals were observed during NREM sleep (Fig. 1E and Extended Data Fig. 1B). We then examined the brains for signs of damage and neuroinflammation by DAPI (Fig. 1H), GFAP (Fig. 1I), and silver staining (Fig. 2G). Except for a minor increase in the number of GFAP-stained astrocytes in deep cortical layers situated directly under the LED (arrows in Fig. 1I and 1K), we did not find any significant differences in these stainings compared to control mice unexposed to red light. Thus, red light induced gliosis in a small area of the brain that did not affect EEG power or sleep.

**Fig. 2.**
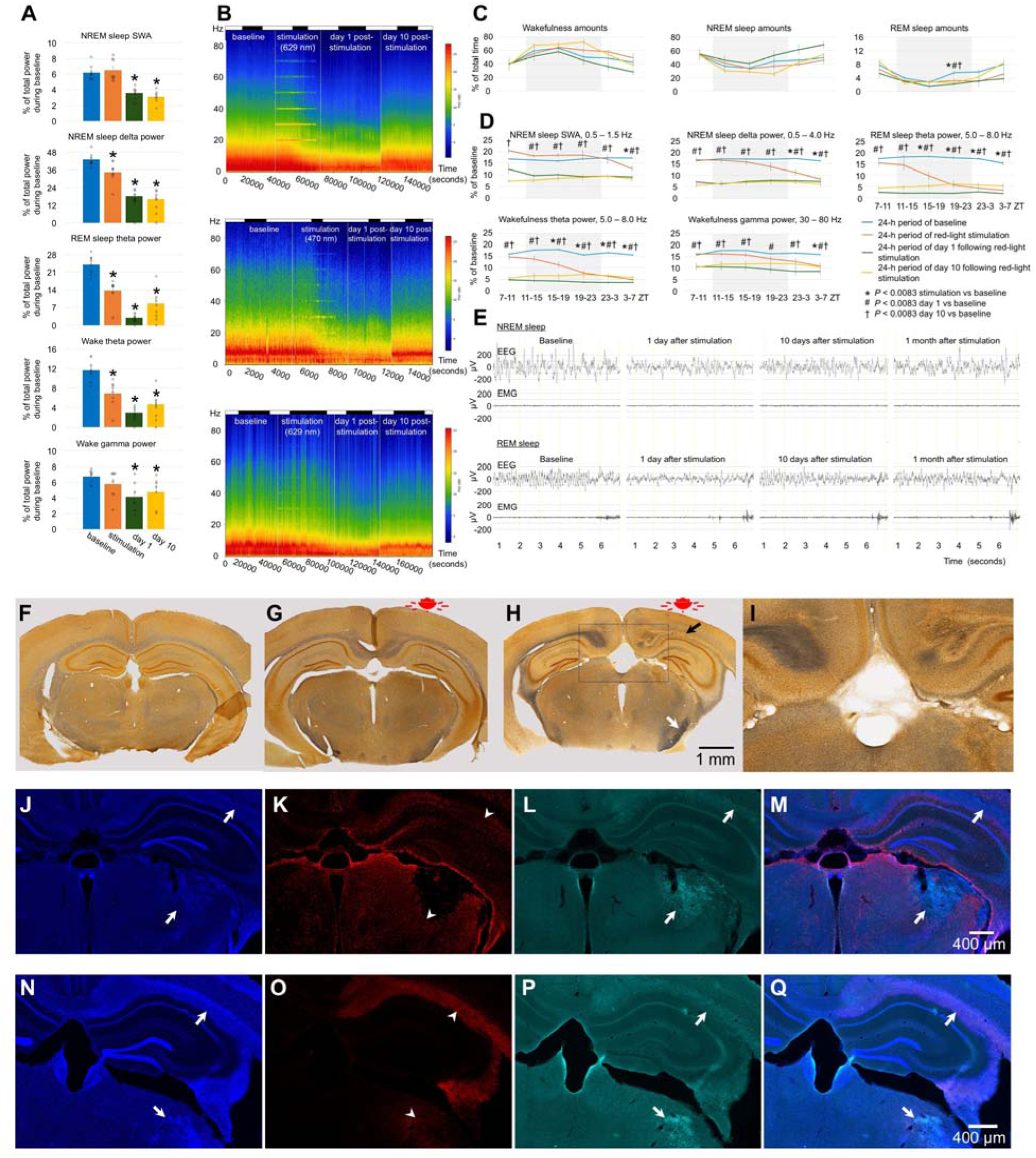
Effect of red light on the brain of older mice. (**A**) EEG power in SWA, delta, theta and gamma ranges. One-way repeated measures ANOVA followed by Tukey’s post hoc test, \**P* < 0.05 vs baseline. (**B**) Representative heatmap of EEG power spectrum during NREM sleep (Top), REM sleep (Middle), and wakefulness (Bottom). (**C**) Amounts of sleep and wakefulness and (**D**) profile of changes of EEG power in SWA, delta, theta and gamma range. Each datapoint represents power calculated during a 4-hour period. One-way repeated measures ANOVA followed by Tukey’s post hoc test, *#†*P* < 0.0083 vs baseline. (**E**) Representative EEG and EMG recordings of NREM sleep (upper panel) and REM sleep (lower panel) during baseline, stimulation, and post-stimulation days in mice exposed to red light. Silver staining of a brain section in a control mouse unexposed to light (**F**), a 3-month-old mouse exposed to red light (**G**), and a 16-month-old mouse exposed to red light (**H** and **I**). Scale bar in H also applies to F and G. (**J**) DAPI staining, (**K**) GFAP staining, (**L**) autofluorescence under a green filter, and (**M**) merged image of J-L in a mouse exposed to red light. (**N**) DAPI staining, (**O**) IBA-1 staining, (**P**) autofluorescence under a green filter, and (**Q**) merged image of N-P in a mouse exposed to blue light. Error bars indicate the standard error of the mean. *P<0.05 vs baseline.

Exposure to blue light caused local damage to the cerebral cortex (Fig. 1L, 1M, 1N, and 1O). The lesioned area displayed autofluorescence (Fig. 1L and 1N) and was surrounded by GFAP-positive astrocytes (arrows in Fig. 1M and 1O). Despite the local brain damage, the amount of wakefulness and sleep did not change significantly (Fig. 1C). EEG powers were reduced across all tested frequency ranges by 20-40% during the stimulation and post-stimulation days (Fig. 1F, 1G, Extended Data Fig. 2A and 2B). Thus, blue light produced local damage in the cerebral cortex and moderately reduced EEG power.

### Effect of blue vs red light on older mice

We then compared the effects of red and blue light irradiation on the brains of older animals. 16-month-old C57BL/6 mice were exposed to red light for 8.5 hours. Wakefulness and sleep amounts did not change significantly except a minor reduction of REM sleep at the end of light-off period (Fig. 2C). Following 6 hours of light stimulation, we observed a large reduction in the EEG powers (40-90% reduction) across all vigilance stages in all monitored frequencies (Fig. 2A, 2B and 2D). Notably, we observed significant reductions in powers within the theta range, which was nearly abolished during REM sleep (Fig. 2A, 2E, and Extended Data Fig. 3).

Interestingly, the power continued to decrease after the light stimulation ended and reached the lowest level at the end of the post-stimulation day (Fig. 2D). In addition to the massive changes in the EEG power, we also observed changes in the animal’s behavior. At the end of the stimulation period and shortly after it, the mice displayed hyperactivity and circling behavior that was followed by hypoactivity. Silver staining indicated extensive damage of the fiber tracts.

Contrary to the brains of mice unexposed to light (Fig. 2F) or young mice exposed to red light (Fig. 2G), both the medial and lateral forebrain bundle systems were damaged in the older animals after red light exposure (Fig. 2H and 2I). Fiber tracts were destroyed directly below the LED so that the silver staining was lacking in this area (black arrow in Fig. 2H). However, they were intensely stained in black color in various other brain areas (Extended Data Fig. 4), even those located distantly from the LED including the ventral parts of the brain (e.g., the cerebral peduncle, white arrow in Fig. 2H). Such dark staining of the fiber bundles was not found in the brains of young mice exposed to red light, suggesting that the fiber damage was prevalent only in the older mice. We also observed autofluorescence along fiber bundles (the cingulum bundle, alveus, corpus callosum, etc.) and within thalamic structures (arrows in Fig. 2J - 2Q, Extended Data Fig. 5). IBA1 immunofluorescence was seen within these locations (arrowheads in Fig. 2O), and GFAP immunofluorescence surrounded them (arrowheads in Fig. 2K). These results suggest the presence of reactive microglia as part of neuroinflammation within the damaged brain structures as well as gliosis in the surrounding areas. Despite the damage to subcortical structures, no visible loss of neurons was detected in the cerebral cortex directly under the LED (Extended Data Fig. 6).

14-month-old mice exposed to blue light for 14.7 hours did not exhibit significant changes in sleep amounts (Fig. 3C). Their EEG power was reduced across a wide range of frequencies during stimulation and/or post-stimulation (Fig. 3A, 3B, Extended Data Fig. 7A and 7B). Exposure to blue light caused the obliteration of tissue in the cerebral cortex (Fig. 3D). The damage spread deeper into the brain because we observed darker silver staining in the internal capsule and nearby areas (arrow in Fig. 3E) as well as the presence of GFAP-positive astrocytes 1.5 mm or more below the LED (arrow in Fig. 3G). The IBA1-positive microglia were absent (Fig. 3K), suggesting a moderate neuroinflammatory response. Compared to red light, blue light ablated cortical tissue locally but produced less damage to the fiber bundles subcortically. EEG power deficits were massive following exposure to red light and moderate following exposure to blue light in older animals.

**Fig. 3.**
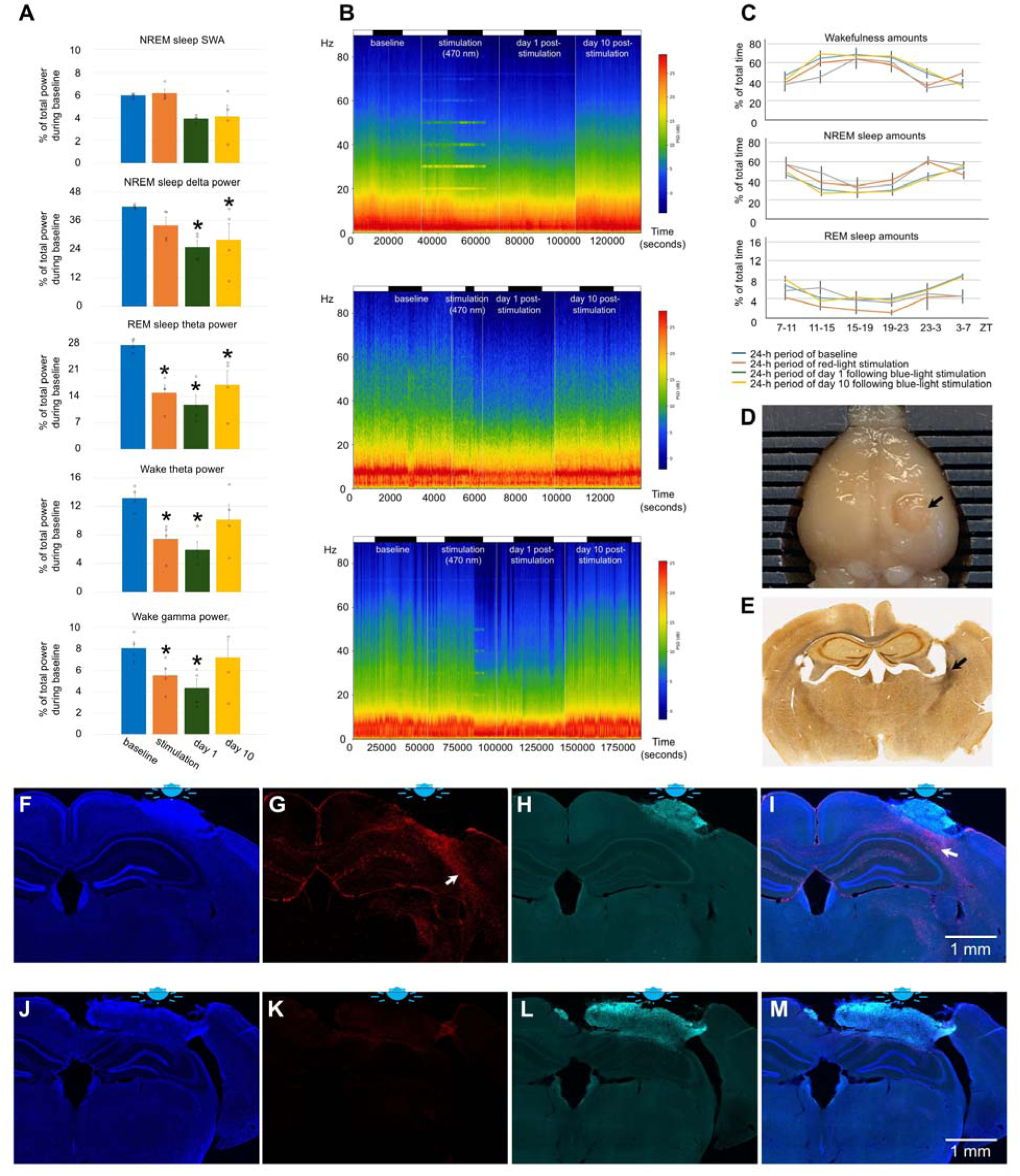
Effect of blue light on the brain of older mice. (**A**) EEG power in SWA, delta, theta and gamma ranges during baseline, stimulation and post-stimulation days in mice exposed to blue light. One-way repeated measures ANOVA followed by Tukey’s post hoc test, \**P* < 0.05 vs baseline. (**B**) Representative heatmap of EEG power spectrum during NREM sleep (Top), REM sleep (Middle), and wakefulness (Bottom) calculated in a mouse exposed to blue light. A bar on the top of the figures indicates light-on and light-off periods. (**C**) Wakefulness (Top), NREM sleep (Middle) and REM sleep amounts (Bottom) in mice exposed to blue light. (**D**) Local damage of the brain under the LED in a mouse exposed to blue light. The damaged area is indicated by the black arrow. (**E**) Silver staining of a brain section of a mouse exposed to blue light. The black arrow indicates intensely stained fiber bundles. (**F**) DAPI staining, (**G**) GFAP staining, (**H**) autofluorescence under a green filter, and (**I**) merged image of F-I in a mouse exposed to red light. (**J**) DAPI staining, (**K**) GFAP staining, (**L**) autofluorescence under a green filter, and (**M**) merged image of J-M in a mouse exposed to blue light. Error bars indicate the standard error of the mean.

### Age- and dose-dependent effect of red light

We tested the effect of red light in two additional age groups, 7.5-month and 23-month-old mice. The average duration of the stimulation was 8.2 ± 0.2 hours in 23-month-old mice and ranged between 6.7 hours and 13.4 hours in 7.5-month-old mice.

Red light exposure caused a slight reduction in wakefulness (Fig. 4A) and moderately consolidated sleep by increasing the bout duration while decreasing the bout number (Extended Data Fig. 8A and 8B) in 7.5-month-old mice. We observed a power reduction in all assessed frequency ranges (Fig. 4C). The reduction was particularly strong in the theta range.

**Fig. 4.**
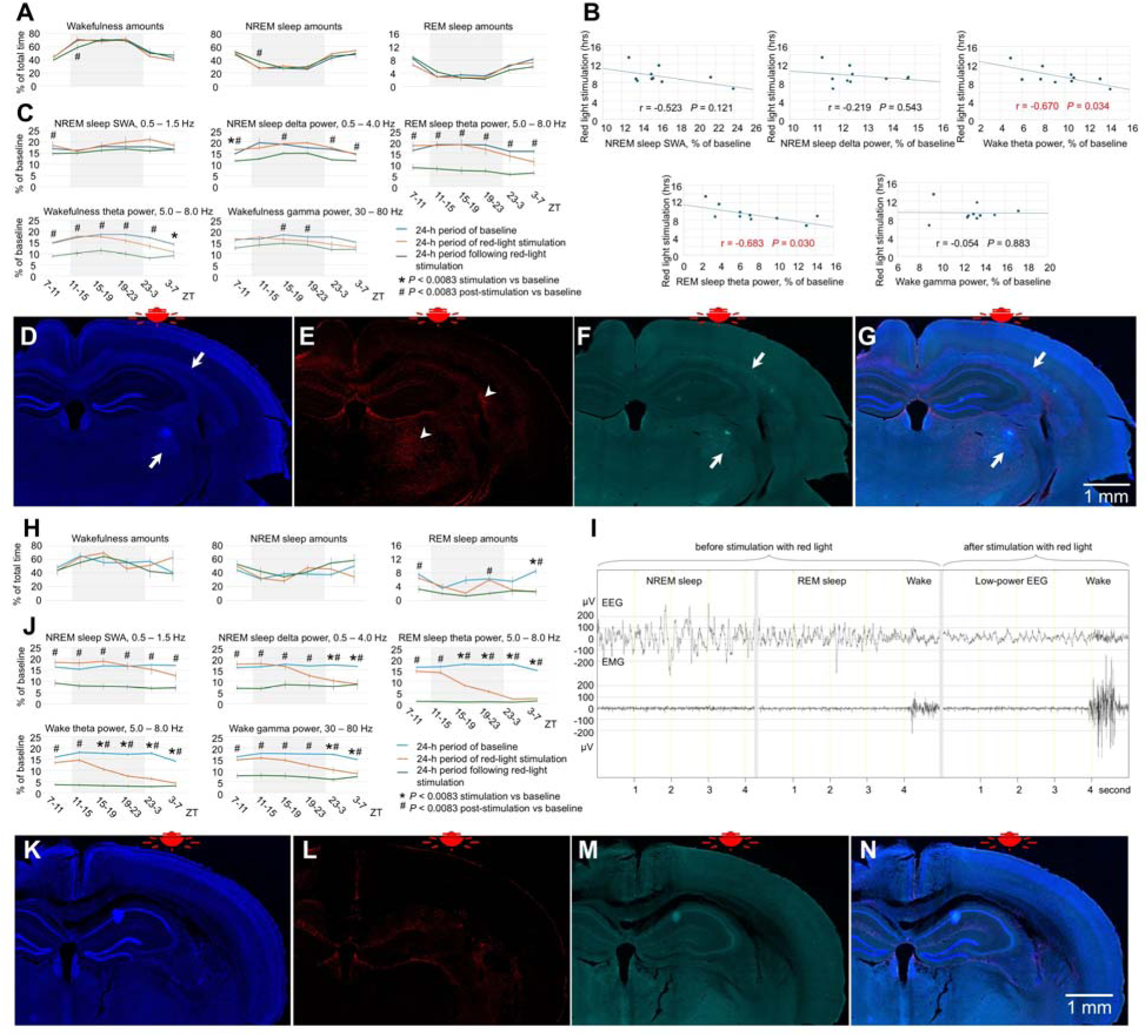
Effect of red light on the brain of 7.5-month-old and 23-month-old mice. (**A**) Amounts of sleep and wakefulness in 7.5-month-old mice exposed to red light. (**B**) Correlation between duration of exposure to red light and EEG power in SWA, delta, theta and gamma ranges in 7.5-month-old mice. (**C**) EEG power in SWA, delta, theta and gamma ranges during baseline, stimulation and post-stimulation days in 7.5-month-old mice exposed to red light. (**D**) DAPI staining, (**E**) GFAP staining, (**F**) autofluorescence under a green filter, and (**G**) merged image of D-F in a 7.5-month-old mouse exposed to red light. White arrows show autofluorescence. (**H**) Amounts of sleep and wakefulness during baseline (blue line), stimulation (orange line) and post-stimulation (green line) days in 24-month-old mice exposed to red light. (**I**) Representative EEG and EMG recordings in a 24-month-old mouse during baseline and post-stimulation day. (**J**) EEG power in SWA, delta, theta and gamma ranges during baseline, stimulation and post-stimulation days in 24-month-old mice. (**K**) DAPI staining, (L) GFAP staining, (M) autofluorescence under a green filter, and (**N**) merged image of K-N in a 24-month-old mouse exposed to blue light. Error bars indicate the standard error of the mean. (**A**, **C**, **H** and **J**) One-way repeated measures ANOVA followed by Tukey’s post hoc test. *#*P* < 0.0083 vs baseline.

Furthermore, the duration of the stimulation negatively correlated with the REM sleep and wakefulness theta power, indicating the dose-dependent response (Fig. 4B). The brain damage was minor. A small enhancement of GFAP immunostaining (arrowheads in Fig. 4E) and autofluorescence within a restricted area of fiber bundles and thalamus (arrows in Fig. 4D, 4F and 4G) were observed. Thus, red light stimulation strongly affected EEG power and caused minor changes in sleep patterns of 7.5-month-old mice.

Despite the duration of red-light stimulation being lower in 23-month-old mice than in 16-month-old mice (8.2 hours vs 8.5 hours), changes in sleep, EEG power and animal behavior were more severe. We observed strong hyperactivity and circling behavior at the end of red-light stimulation, which was followed by immobility. Hence, three of the mice could not eat and drink and had to be euthanized. Theta power was nearly abolished during REM sleep and wakefulness. Large reductions in NREM sleep delta power and wakefulness gamma power were also observed (Fig. 4J). At a later stage when the mice became immobile, the EEG displayed a regular pattern of low-voltage and low-frequency activity that was sometimes interrupted by brief episodes of increased EMG and desynchronized EEG (Fig. 4I). It was not possible to recognize either REM or NREM sleep during this stage based on the EEG pattern. We analyzed the extent of brain damage in 23-month-old mice by GFAP (Fig. 4L) and IBA immunostaining as well as autofluorescence (Fig. 4M). Interestingly, these markers were not detected in the brains of 23-month-old mice that were euthanized within 24 hours of stimulation offset. The likely explanation is an insufficient period (> 24 hours) for these markers to develop.

### Longer red-light exposure at lower intensity

The lowering red light intensity by ∼3-fold and increasing the light beam diameter by ∼1.5-fold was achieved by placing a round piece of aluminum foil directly under the LED (Fig. 5A and 5B). After 3 days of brain exposure of 15-month-old mice to red light (8.5 hours per day), the mice displayed a rapid reductions of EEG power across all assessed frequency ranges, with REM sleep theta power nearly abolished (Fig. 5C and 5E). Wakefulness amounts decreased and NREM sleep amounts increased during day 4 of red-light stimulation and post-stimulation days, returning to the original levels on days 10 and 30 post-stimulation (Fig. 5D and 5F). We then compared the NREM sleep spindle density between 3-month-old and 16-month-old mice exposed to 8.5 hours of red light at a higher intensity and 15-month-old mice exposed to red light for 34 hours at a lower intensity. The spindle density did not change in young mice but was markedly reduced in older mice (Extended Data Fig. 9). The histological analysis (Extended Data Fig. 10) revealed that the longer red-light exposure at a lower intensity produced similar damage to the brain as that found in 16-month-old mice exposed to 8.5 hours of red light at a higher intensity.

**Fig. 5.**
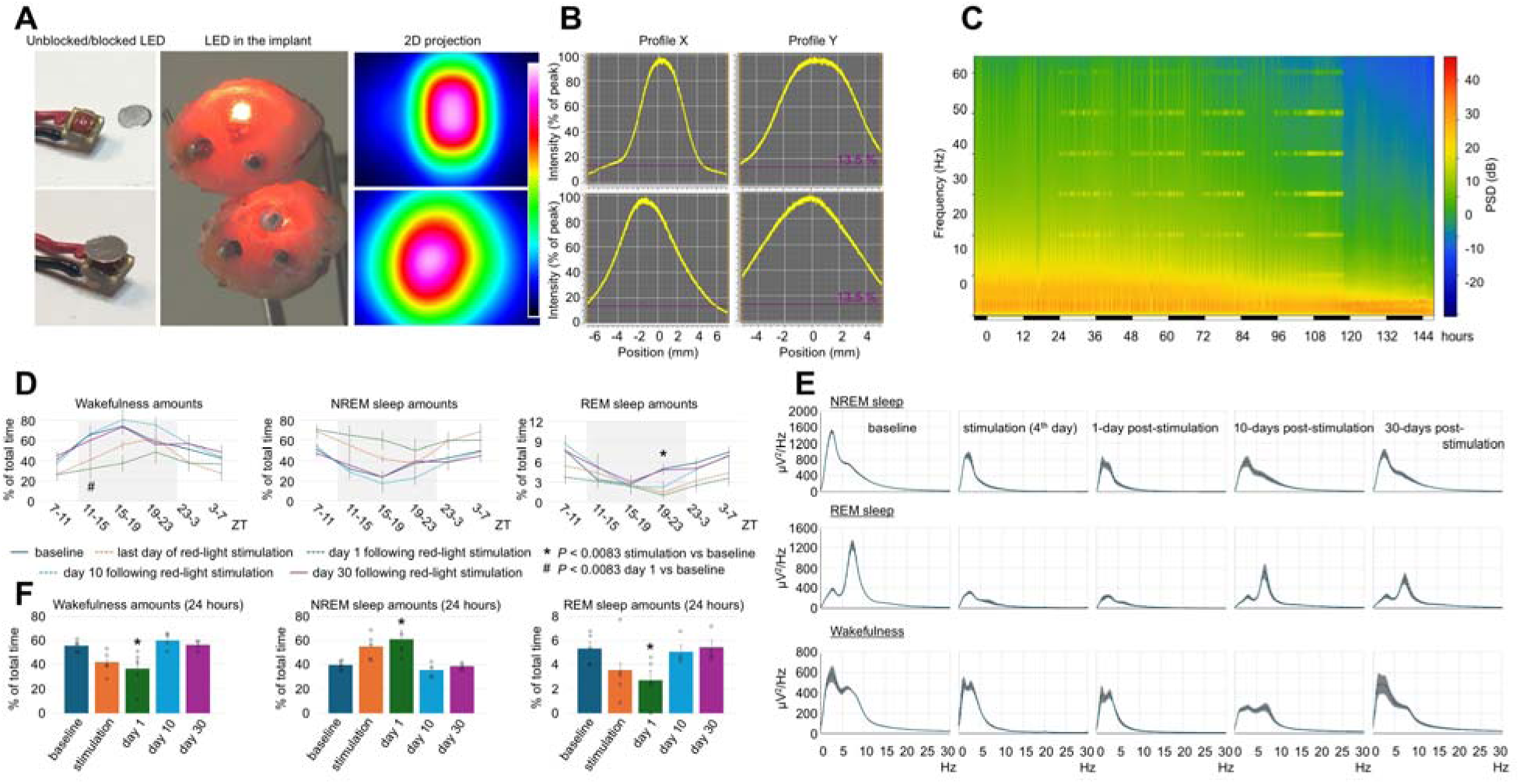
Effect of red light delivered at lower power and longer duration on the brain of 15-month-old mice. (**A**) The unblocked and blocked LED (Left), its location within the head implant (Middle), and 2d projection of the light beam produced by the unblocked and blocked LED (Right). (**B**) Profiles of beam intensity of unblocked (Top) and blocked (Bottom) LEDs. (**C**) Representative heatmap of EEG power spectrum during NREM sleep recorded in a mouse during baseline, day 4 of red light of stimulation, and post-stimulation day. (**D**) Profile of changes of wakefulness, NREM and REM sleep in mice during the last day of red-light stimulation. Each datapoint represents amounts of wakefulness or sleep calculated during a 4-hour period. One-way repeated measures ANOVA followed by Tukey’s post hoc test. *#*P* < 0.0083 vs baseline. (**E**) Power spectral density of the EEG during NREM sleep (Top), REM sleep (Middle) and wakefulness (Bottom) during baseline, day 4 of red light of stimulation, and post-stimulation days. (**F**) 24-hour amounts of wakefulness, NREM and REM sleep were calculated in mice on day 4 of red-light stimulation. Error bars indicate the standard error of the mean. \**P*<0.05 vs baseline.

## Discussion

Blue light is considered significantly more damaging to brain cells than red light due to its shorter wavelength and higher energy^3^. Consistent with high phototoxic properties of light with shorter wavelength, 8.5-hour exposure to blue light resulted in extensive local damage of neurons under the LED in 3-month and 14-month-old mice. Sleep amounts did not change significantly, and EEG power was moderately reduced in these mice.

Red light, with a longer wavelength and lower energy, is considered less phototoxic and is used in photobiomodulation therapy^13, 14^. Consistent with low phototoxic properties, red light did not produce any noticeable damage to the neurons and did not change sleep amounts significantly in young mice. Surprisingly, brain exposure to red light had a strong effect on older animals. While sleep duration was not significantly affected, EEG power was reduced massively, especially in the theta range during both stimulation and post-stimulation periods in 16-month-old mice. This reduction in EEG power was likely caused by damage to fiber tracts throughout the brain. Several fiber bundles have been destroyed at least partially under the LED, which could explain the reduction of delta power (the corpus callosum, external capsule, and cingulum bundle) and theta power (the alveus)^15, 16^. In addition, damage to the fimbria could contribute to the massive reduction in theta power^17^, whereas the damage to the internal capsule could explain the immobility seen in the mice during the post-stimulation day^18^. The effect of red light on the brain of 23-month-old mice was even more severe. Three of 7 mice in this group entered a coma-like state during which they could not eat and drink and therefore had to be euthanized.

Red light likely produced damage more severe compared to blue light to the brain of old mice because it can penetrate deeper into the brain tissue^19–21^. The damage depended on the type of tissue exposed to light. We observed loss of fibers and tissue lesions in the thalamus 2 mm or more from the brain surface. The light power density was expected to be up to 100-fold higher in the cerebral cortex directly under the LED compared to deeper brain areas^19–21^, yet no visible neuronal loss was seen in the cerebral cortex. Furthermore, none of the sleep regulatory areas must have been significantly damaged by red light exposure since minor changes in amounts of wakefulness and sleep were found in our studies. The differences in the effects of red light on EEG power and sleep were particularly evident in 7.5-month-old mice, in which massive changes in EEG power that correlated with the duration of the stimulation and minor changes in sleep pattern were observed.

Light may have either damaging or therapeutic properties depending on the power density and duration of the treatment^22, 23^. These parameters were higher in our experiments than in photobiomodulation therapies^24^. Nevertheless, uncontrolled use of red-light LED helmets and panels that are freely available on the market to humans could potentially lead to high brain exposure to light, approaching the levels used in our study. Moreover, pulsed delivery of red light has been considered in photobiomodulation therapy^25–27^. It is important not to overexpose the brain of older people to light because fiber bundles appear especially vulnerable to pulsed red-light exposure. Additionally, it is unclear what irradiation level of red light would be safe if treatment were done over extended periods. These levels need to be empirically determined and tested. We reduced the power density of red light by approximately 3-fold by placing a round piece of aluminum foil under the LED. When delivered over 4 days (34 hours of light stimulation), red light produced a similar reduction in EEG power as that seen in the mice treated with higher irradiation for 1 day (8.5 hours of light stimulation), suggesting that light exposure is additive over time. Additional studies are needed to assess the irradiation and fluence levels at which red light does not cause any side effects. This is important because high levels of fluence can be achieved even during exposure to sunlight if it occurs over a period of months or years. Recent studies demonstrated a positive correlation between outdoor light time and incident dementia^28^, and between prolonged sunlight exposure and structural changes in the brain^29^. The possible role of sunlight penetrating through the skull and contributing to these pathologies cannot be ruled out.

Earlier studies demonstrated that the damaging effects of blue light in the Drosophila model were age dependent^11^. Blue light exposure was associated with reduced activity of specific components of energy-producing pathways in mitochondria^30^. Mitochondrial dysfunction has been associated with aging in mammals and in variety of neurodegenerative diseases^31, 32^. Therefore, it is a likely mechanism responsible for the brain damage associated with red light in older animals. However, additional factors may contribute to the effects of red light on an aging brain, including oxidative stress, changes in microcirculation and accumulation of lipofuscin in aged brain cells^33–35^. Future studies are necessary to define the exact mechanism(s) involved.

## Supporting information

The Excel file contains original data of the study.

## Methods

### Animals

C57BL/6J male mice were obtained from the Jackson Laboratory (stock # 000664) or bread in house at the vivarium of West Roxbury VA, Massachusetts, USA. Mice were housed at 70 - 74°F and a 12-h light/dark cycle (7:00 AM–7:00 PM) with food and water ad libitum. All experiments conformed to US Veterans Administration, Harvard University, and US National Institutes of Health guidelines.

### Experimental groups

The performed study included 6 experimental groups. The number, sex, and age of the animals in each experiment were following.

Experiment 1. Effect of red light on brains of young mice

n=9, 2 females and 7 males, 2.79 ± 0.05 month-old, Fig. 1B, 1D, 1E, 1H–1K, Extended Data Fig. 1

Experiment 2. Effect of blue light on brains of young mice

n=5, 1 female and 4 males, 3.09 ± 0.00 month-old, Fig. 1C, 1F, 1G, 1L–1O, Extended Data Fig. 2

Experiment 3. Effect of red light on brains of older mice

n=8, 4 females and 4 males, 16.08 ± 0.07 month-old, Fig. 2, Extended Data Fig. 3 and 4

Experiment 4. Effect of blue light on brains of older mice

n=4, 2 females and 2 males, 13.73 ± 0.27 month-old, Fig. 3, Extended Data Fig. 7

Experiment 5. Dose-dependent effect of red light

n=10, 2 females and 8 males, 7.54 ± 0.09 month-old, Fig. 4A-4G, Extended Data Fig. 8

Experiment 6. Effect of red light on brains of 23-month-old mice n=7, 3 females and 4 males, 23.11 ± 0.51 month-old, Fig. 4H-4N

Experiment 6. Effect of red light on the brain delivered at a lower intensity for a longer time

n=5, males, 14.96 ± 0.00 month-old, Fig. 5, Extended Data Fig. 10

Additionally, 4 untreated C57BL/6 mice, 3-6 month-old, were used as a control in silver staining studies.

### Stereotaxic surgery

Mice were deeply anesthetized with isoflurane (1–3%) and fixed to a stereotaxic frame. The skin over the skull was incised and retracted to expose the skull. Electroencephalogram (EEG) electrode was screwed into the skull above the parietal cortex (anteroposterior, -2.0 mm; lateral, 2 mm over left hemisphere of the brain), and a reference electrode screwed into the skull above the cerebellum. Electromyography (EMG) electrodes were placed in the nuchal muscle. A custom-made LED light source (629 nm, L1C1-RED1000000000 or 475 nm, L1C1-BLU1000000000, Lumileds) was placed on the skull above the right parietal cortex on the contralateral side to EEG electrodes and fixed in place with uncolored dental cement (Stoelting) for subsequent light stimulation. The electrodes/LED were connected to a plug (PlasticsOne) and fixed to the skull using Fusion Flo flowable composite (Prevest Denpro). The scalp incision was sutured closed. The mice were injected with meloxicam and allowed to recover.

### In vivo EEG/EMG recordings

Recordings were performed in freely moving animals following 2 days of acclimatization to the recording chamber. After acclimatization, the wires from the plug were connected to a free-moving container close to the mouse (Neuroratgeting Systems) that housed the three-channel transmitter (Pinnacle Technology’s 8200-K9-SL, an EEG/EMG wireless mouse system for sleep). For recording EEG/EMG, the USB dongle receiver was connected to the computer. The Sirenia acquisition software was used to access the USB dongle (Part #8406-SL, Pinnacle Technology, Inc., Lawrence, KS), and the data were sampled at 512 Hz. EEG and EMG signals were acquired, digitized, and amplified by the transmitter before being sent via Bluetooth to the USB receiver. EEG data were sampled using the parietal and reference electrode (left posterior screw), and the EMG signal was generated via the difference between the two wire leads in the nuchal muscles.

### Exposure of mouse brain to either blue or red light

Adult C57BL/6J mice were subjected to either red light (629 nm) or blue light (475 nm) stimulation. In most mice, EEG/EMG recordings were performed continuously from day 4 (day 4 is the baseline day, day 5 is the stimulation day, day 6 is the post-stimulation day, and day 15 is the post-stimulation 10th-day recovery) up to day 15. In some mice, recordings were performed for more than a month, whereas some mice had to be euthanized during the post-stimulation day due to immobility. The light stimulation by either red or blue light was given at 10 Hz during online detection of the NREM sleep state, which mainly occurred within the 24-hour recording period. Light stimulation began starting at ZT7.

### Offline sleep data analysis

The data file saved in Sirenia software was imported into SleepSign software (Kissei Comtec) to score sleep-wake states in 5-sec epochs further. Sleep scoring during the 24-h recording on day 4 (baseline day) was compared to light-stimulation day (day 5) and also to day 6 of the post-stimulation day, and day 15 of post-stimulation 10th day recovery in young and old mice exposed to red (629 nm) or blue (475 nm) light. The behavioral states, namely non-rapid eye movement (NREM) sleep, REM sleep, and waking, were analyzed in a 4-hr time bin as previously described^36^ and determined based on the following criteria: (1) Wake; active behavior accompanied by desynchronized EEG (low in amplitude), and tonic/phasic motor activity evident in the EMG signal; (2) NREM sleep; more synchronized EEG, higher in amplitude, with particularly notable power in the delta (0.5-4.0 Hz) band, and low motor activity (EMG); and (3) REM sleep; small amplitude EEG, particularly notable power in the theta (5-8 Hz) band, and phasic motor activity (EMG). Compared to wake, EEG power during REM sleep was significantly reduced in delta frequencies (0.5-4.0 Hz) and increased in the range of theta activity (5-8 Hz). The measures included calculating the amount and % time spent in the wake, NREM sleep or REM sleep of the 24-h recording during the baseline day and the data compared to light stimulation day (day 5) vs post-stimulation day (day 6) vs 10th-day post-stimulation (day15) recovery day in young and old mice. The average duration and number of bouts for each state were calculated as indices of the consolidation of behavioral states.

### Spectral analysis of EEG signal

To analyze the EEG data, multitaper spectral analysis (Chronux Toolbox; http://chronux.org) was performed on 24 h raw EEG data, using custom scripts written for MATLAB (R2016a-2018b, MathWorks) as described previously^36, 37^. The multitaper method works by averaging together multiple independent spectra estimated from a single segment of data, which has been shown to have superior statistical properties compared with single-taper spectral estimates^38^. Power spectral density for each 5-s EEG epoch was computed for each sleep-wake state using a Fast Fourier transform (FFT) after using a Hanning window, yielding power density spectra with a frequency resolution of 0.12207 Hz. Epochs containing artifacts (more than 10xSEM) were excluded from spectral analysis. To account for inter-individual differences in absolute EEG power spectra, power spectra density (PSD) in each frequency 4-h bin and for each state was expressed as a percentage of a reference value, calculated across the 24-h recording in baseline day, stimulation day, post-stimulation day and post-stimulation 10th-day recovery (and 30th-day post-stimulation recovery day for chronic light stimulation), for each mouse as the mean total EEG power in each frequency 4-h bins [slow wave activity (0.5-1.5 Hz), delta (0.5–4.0 Hz), theta 5.0-8.0 Hz, and gamma (30.0-80.0 Hz)] for each of the behavioral states (NREM sleep or wake or REM sleep states)^36^. The total power for each investigation (NREM sleep or wake or REM sleep states) was then averaged across all mice. For statistical analysis of this data, absolute power across the four frequency bands of interest [slow wave activity (0.5-1.5 Hz), delta (0.5–4.0 Hz), theta 5.0 -8.0 Hz and gamma (30.0-80.0 Hz)] for each 24 h EEG segment for NREM, REM sleep and wake values were compared between baseline vs light-stimulation vs post-stimulation vs post-stimulation recovery recording using one-way analysis of variance or a repeated measure analysis of variance followed up by a Tukey’s multiple comparisons test. Differences were determined to be significant when P<0.05.

Power in each frequency band, namely SWA (0.5-1.5 Hz), delta (0.5-4.0 Hz), theta (5.0-8.0 Hz) and Gamma 30.0-80.0 Hz), was calculated by averaging the power values obtained from the power spectral densities in the corresponding frequency bands, excluding 60 Hz (line noise).

### Power spectral density heatmaps and plots

To generate these heatmaps and plots, we used Python implementation of the script developed in Prerau Lab^37^. EEG datasets were processed by this script separately for wakefulness, NREM sleep and REM sleep using the following parameters.

fs = 512 # Sampling Frequency

frequency_range = [0, 90] # Limit frequencies from 0 to 90 Hz

time_bandwidth = 3 # Set time-half bandwidth

num_tapers = 5 # Set number of tapers

window_params = [5, 5] # Window size is 5s with step size of 5s

min_nfft = 0 # No minimum nfft

detrend_opt = ‘constant’ # detrend each window by subtracting the average

multiprocess = True # use multiprocessing

cpus = 3 # use 3 cores in multiprocessing

weighting = ‘unity’ # weight each taper at 1

plot_on = True # plot spectrogram

return_fig = False # do not return plotted spectrogram

clim_scale = True # do not auto-scale colormap

verbose = True # print extra info

xyflip = False # do not transpose spect output matrix

This program generated both heatmaps and spectrogram output matrices in the range of 0-90 Hz and frequency bin of 0.125 Hz. The matrices were processed to remove outliers that were 10x or more of the standard deviation in each frequency bin and then used to generate power spectral density plots.

### Spindle detection

Sleep spindles were detected using an automated algorithm (MATLAB) validated and published earlier^39, 40^. Briefly, EEG data was band-pass filtered (10.0–15.0 Hz, Butterworth Filter), and the root-mean-squared (RMS) power was calculated to provide an upper envelope of the data. The RMS data was then exponentially transformed to further accentuate spindle-generated signals over baseline. Putative spindle peaks were identified in transformed data via crossing of an upper-threshold value, set as 3.5x the mean RMS EEG power across all states for each mouse. Additional detection criteria included a minimum duration of 0.5 s, based on the crossing of a lower threshold set at 1.2x mean RMS power, and a minimum inter-event interval of 0.5 s.

### Autofluorescence

Autofluorescence of brain tissue was analyzed in the current study because it is a sign of brain damage. Autofluorescence was seen in a wide range of light spectrum ranging from the blue to green color, although it was particularly strong using Opal 480 imaging filter (excitation: 450 nm, emission: 500 nm, AKOYA Biosciences; Extended Data Fig. 5A and 5B) in the tissue damaged by light. We tested several filters for the presence of autofluorescence in these brain sections and found that it was not visible using Opal 620, 690 and 780 imaging filters (excitation: 588 nm or more, emission: 616 nm or more, AKOYA Biosciences; Extended Data Fig. 5C1 – 5C7 and 5D1 – 5D7). Based on these results, we selected Opal 480 imaging filter to identify the location of tissue damage and used Alexa Fluor 647 label (excitation: 650–653 nm, emission: 665–671 nm) in immunofluorescent studies to avoid interference of the stained markers with autofluorescence.

### Immunohistochemistry

Mice were deeply anesthetized and perfused with 1x phosphate buffered saline (Sigma-Aldrich, catalog # P4417) followed by 10% neutral buffered formalin solution (Sigma-Aldrich, catalog # HT501128). Brains were post-fixed in 10% neutral buffered formalin solution overnight and then placed into 30% sucrose in phosphate buffered saline at 4°C for 24 hours or longer. Brain sections (40 µm thickness) were cut using a freezing microtome (Leica, CM1800) at -20°C and collected in cryoprotective buffer consisting of 1x phosphate buffered saline, 20% glycerol (Sigma-Aldrich, catalog # G5516), and 30% ethylene glycol (Sigma-Aldrich, catalog # 102466). Brain slices were washed in 1x phosphate buffered saline and then blocked using 5% normal donkey serum (Sigma-Aldrich, catalog # S30-M) and 1% Triton-X100 (Sigma-Aldrich, catalog # X100) in 1x phosphate buffered saline for 1 hour at 20°C. Slices were incubated with primary antibodies overnight at 20°C. Brain slices were washed in 1x phosphate buffered saline and incubated in either biotinylated anti-rabbit antibody (Vector Laboratories, catalog # BA-1000) or biotinylated anti-mouse antibody (Vector Laboratories, catalog # BA-9200) for 2-3 hours at 20°C. Brain slices were washed again and then incubated with Alexa Fluor 647 Streptavidin (Jackson ImmunoResearch Laboratories, catalog # 016-600-084). Sections were mounted on Premium Superfrost Plus microscope slides (VWR International, catalog # 48311-703) and cover-slipped with Fluoromount-G mounting medium with DAPI (Thermo Fisher Scientific, catalog # 00-4959-52). In the case when biotinylated anti-mouse antibody was used, brain sections were initially treated with M.O.M. Mouse Ig Blocking Reagent (Vector Laboratories, catalog # MP-2400) according to the manufacture recommendations. The primary antibodies used in this study were rabbit anti-GFAP (D1F4Q) XP^®^ (1:500, Cell Signaling Technology, catalog # 12389), rabbit anti-IBA1 (1:500, Thermo Fisher Scientific, catalog # PA5-21274), and mouse anti-NeuN (1:500, EMD Millipore Corporation, catalog # MAB377). Specificity of the antibodies was confirmed in control immunostainings in which the primary antibody was omitted from the staining procedure.

### Silver staining

Silver staining was performed using the FD NeuroSilver Kit II (FD NeuroTechnologies, Ellicott City, MD) according to the manufacturer’s instructions. The free-floating protocol optimized by the manufacturer (FD NeuroBiotechnologies) was strictly followed, with special precautions to avoid sodium and ethanol. NeuroSilver-stained slices were mounted on Superfrost Plus microscope slides (VWR International, catalog # 48311-703), dried overnight, and cover-slipped with VectaMount Express Mounting Medium (Vector Laboratories, catalog # H-5700-60). Both immunostained- and NeuroSilver-stained slices were scanned using Vectra Polaris™ Automated Quantitative Pathology Imaging System.

### Power density measurements and beam profiling of LED-emitted light

The intensity of constant red light emitted by red LED (629 nm, L1C1-RED1000000000, Lumileds) or blue LED (475 nm, L1C1-BLU1000000000, Lumileds) (100 mW-150 mW) was measured by the PM120D Digital Power & Energy Console (Thorlabs). The power was estimated to be 10-fold lower for the light delivered at the 10 ms on/90 ms off pattern (10% duty cycle). The light beam diameter was calculated from the area that included 86.5% of irradiation emitted by the LED using BC210CV/M CMOS Camera Beam Profiler (Thorlabs).

### Closed-loop stimulation system

The closed-loop stimulation system was used to expose mouse brains specifically during NREM sleep. In the closed-loop system, real-time EEG power spectrum analysis was achieved by acquiring biopotentials from commercial sleep-recording software via user datagram protocol (UDP) communication and powering LEDs implanted on the skull of mice during NREM sleep. \ Powering the LED was done at the 10 ms LED on/90 ms LED off pattern generated by Arduino Uno. It was done by a power bank battery connected to the LED via a resistor and gated by a transistor. Further details of the system have been previously published^12^. The Python script for closed-loop stimulation is freely available from the developers of this software (Neuroratgeting Systems) on request.

### Statistical analysis

Results are reported as means ± standard error. Statistical analysis of EEG power spectra, NREM spindle density and sleep/wake data was performed using SigmaPlot software v12 (San Diego, California, USA)^36^. EEG power spectra were assessed using one two-way repeated-measures analysis of variance (rANOVA) as an omnibus test. We first performed the normality test on each dataset using Shapiro-Wilk test. Parametric tests were used if the dataset pass the normality test. Otherwise, non-parametric tests were used. Kruskal-Wallis rank test was used when the data were not normally distributed. One-way ANOVA followed by Dunn’s method or Tukey’s post hoc test were used for comparisons of three or more treatment groups to determine significant effects. Differences were determined to be significant when P<0.05.

For calculating total sleep-wake amounts between light stimulation treatment vs baseline vs post-stimulation vs post-stimulation 10th day following stimulation, paired comparisons of means were evaluated using one-way ANOVA. Further, statistical analysis was run using statistical analysis software (SigmaPlot v12.3, San Diego, California, USA) and generally entailed a one-way ANOVA with any significant interactions followed up by a Tukey’s multiple mean comparisons test and differences were determined to be significant when P<0.05.

Figures 1B, 1C, 2C, 2D, 3C, 4A, 4C, 4H, 4J, 5D and Extended Data Figures 1A, 2A, 7A included up to 5 groups (baseline, stimulation, post-stimulation, day 10, and day 30) and six four-hour intervals (ZT7-ZT11, ZT11-ZT15, ZT15-ZT19, ZT19-ZT23, ZT23-ZT3, and ZT3-ZT7). To analyze the data in these figures, we initially used either Tukey or Kruskal-Wallis correction for multiple groups. Because we performed six time-interval tests, we divided the original significance level of 0.05 by 6, resulting in a new threshold of 0.0083. Therefore, only p-values less than or equal to 0.0083 were considered statistically significant after adjustment in these figures.

Statistical analysis of NREM spindle density (NREM spindles/min) in each mouse was evaluated during light stimulation vs baseline/post-stimulation recordings and the means from all mice were compared using one-way ANOVA followed by Tukey’s post hoc test and differences were determined to be significant when P<0.05.

## Data availability

The data that support the findings in this study are available in the source data and can be requested from the corresponding authors. Source data are provided with this paper.

## Acknowledgments

We thank S. Gonzalez for assistance with data analysis. This study was supported by National Institutes of Health grant R01AG081809 (DG) and R01AG066171 (KK), Chan Zuckerberg Initiative grant (KK), Cure Alzheimer’s Fund grant (KK).

## Author contributions

D.G., S.T., and K.K. conceived the study. D.G., S.T. designed experiments. D.G., D.A., and S.T. conducted the experiments. D.G., D.A., K.K., S.T. contributed to interpretation of results. D.G. and K.K. contributed to funding acquisition. D.G. and S.T. wrote the manuscript. D.G., D.A., K.K. and S.T. edited the manuscript.

## Competing interests

D.G. is a Research Health Scientist at Veterans Affairs Boston Healthcare System, Boston, MA. The contents of this work do not represent the views of the US Department of Veterans Affairs or the United States Government.

**Correspondence and requests for materials** should be addressed to Dmitry Gerashchenko.

## Extended data

**Extended Data Fig. 1.**
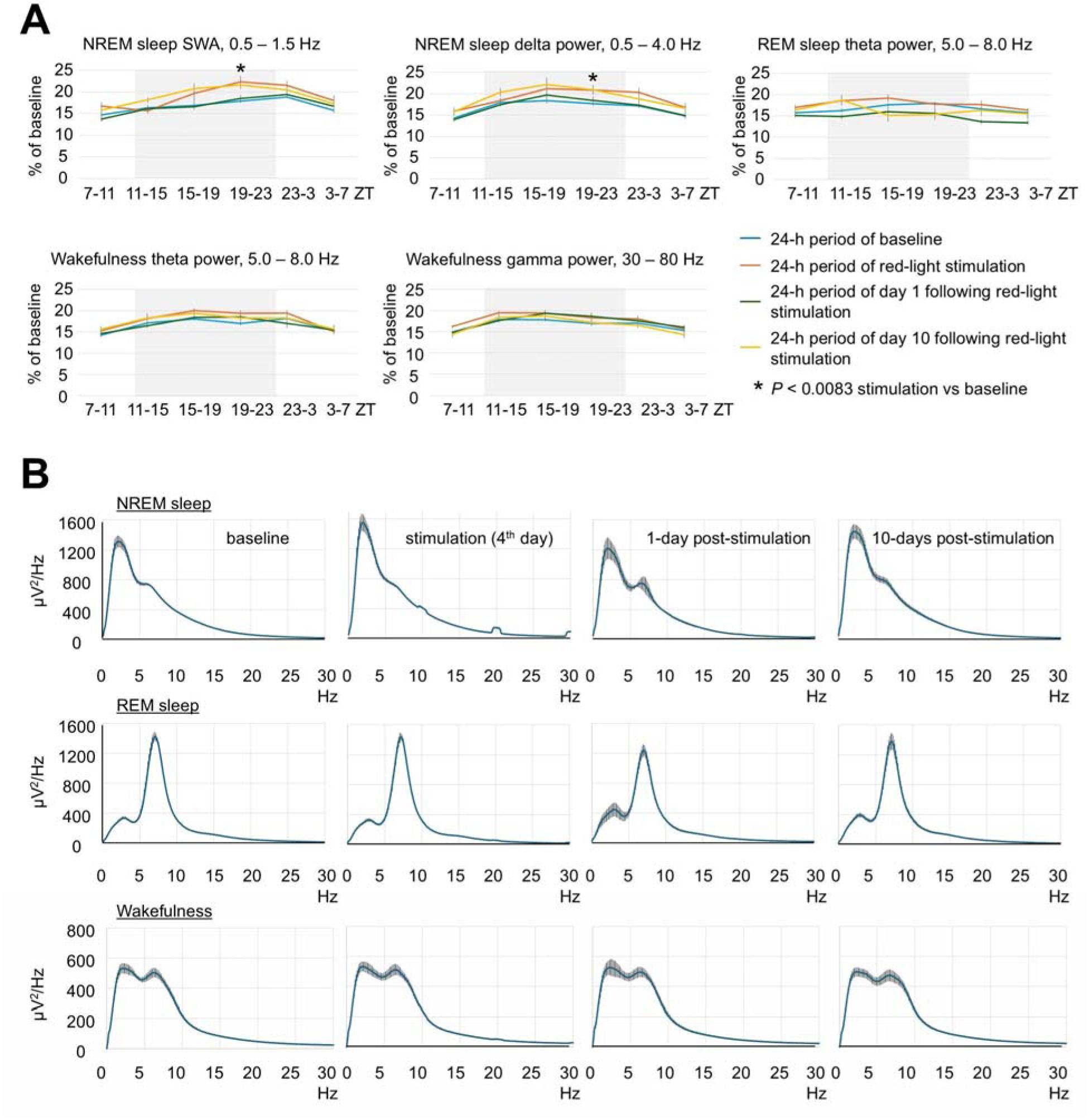
Effect of brain exposure to red light on sleep amounts and EEG power in 3-month-old mice. (**A**) Profile of changes of EEG power in SWA, delta, theta and gamma range in mice exposed to red light. Each datapoint represents power calculated during a 4-hour period. One-way repeated measures ANOVA followed by Tukey’s post hoc test. \**P* < 0.0083 vs baseline. (**B**) Spectral distribution of EEG power density in the frequencies between 0.5 and 30 Hz during NREM sleep (Top), REM sleep (Middle) and wakefulness (Bottom) in mice calculated during baseline, stimulation and post-stimulation days. Error bars indicate the standard error of the mean.

**Extended Data Fig. 2.**
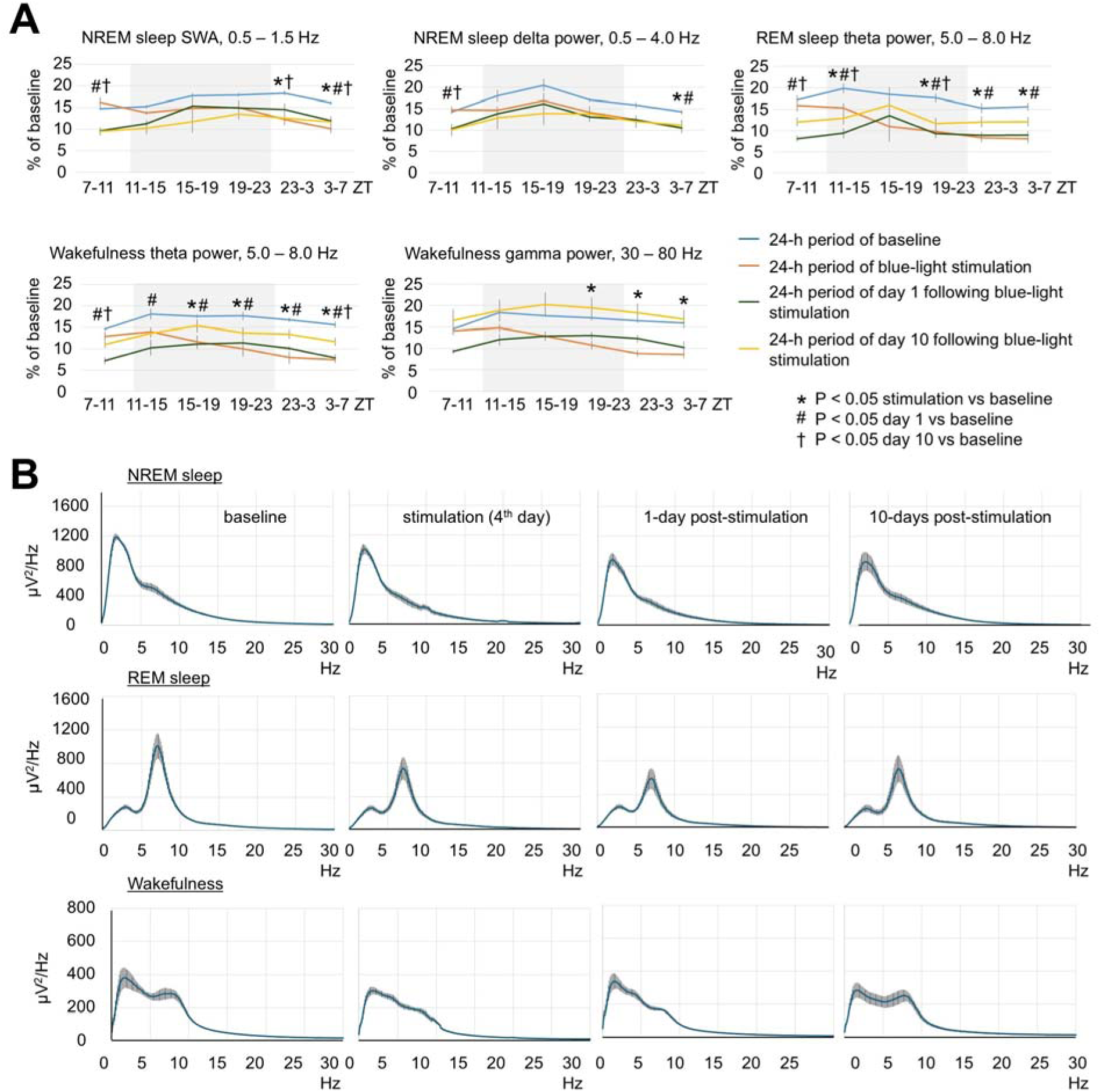
Effect of brain exposure to blue light on sleep amounts and EEG power in 3-month-old mice. (**A**) Profile of changes of EEG power in SWA, delta, theta and gamma range in mice exposed to blue light. Each datapoint represents power calculated during a 4-hour period. One-way repeated measures ANOVA followed by Tukey’s post hoc test, *#†*P* < 0.0083 vs baseline. (**B**) Spectral distribution of EEG power density in the frequencies between 0.5 and 30 Hz during NREM sleep (Top), REM sleep (Middle) and wakefulness (Bottom) in mice calculated during baseline, stimulation and post-stimulation days. Error bars indicate the standard error of the mean.

**Extended Data Fig. 3.**
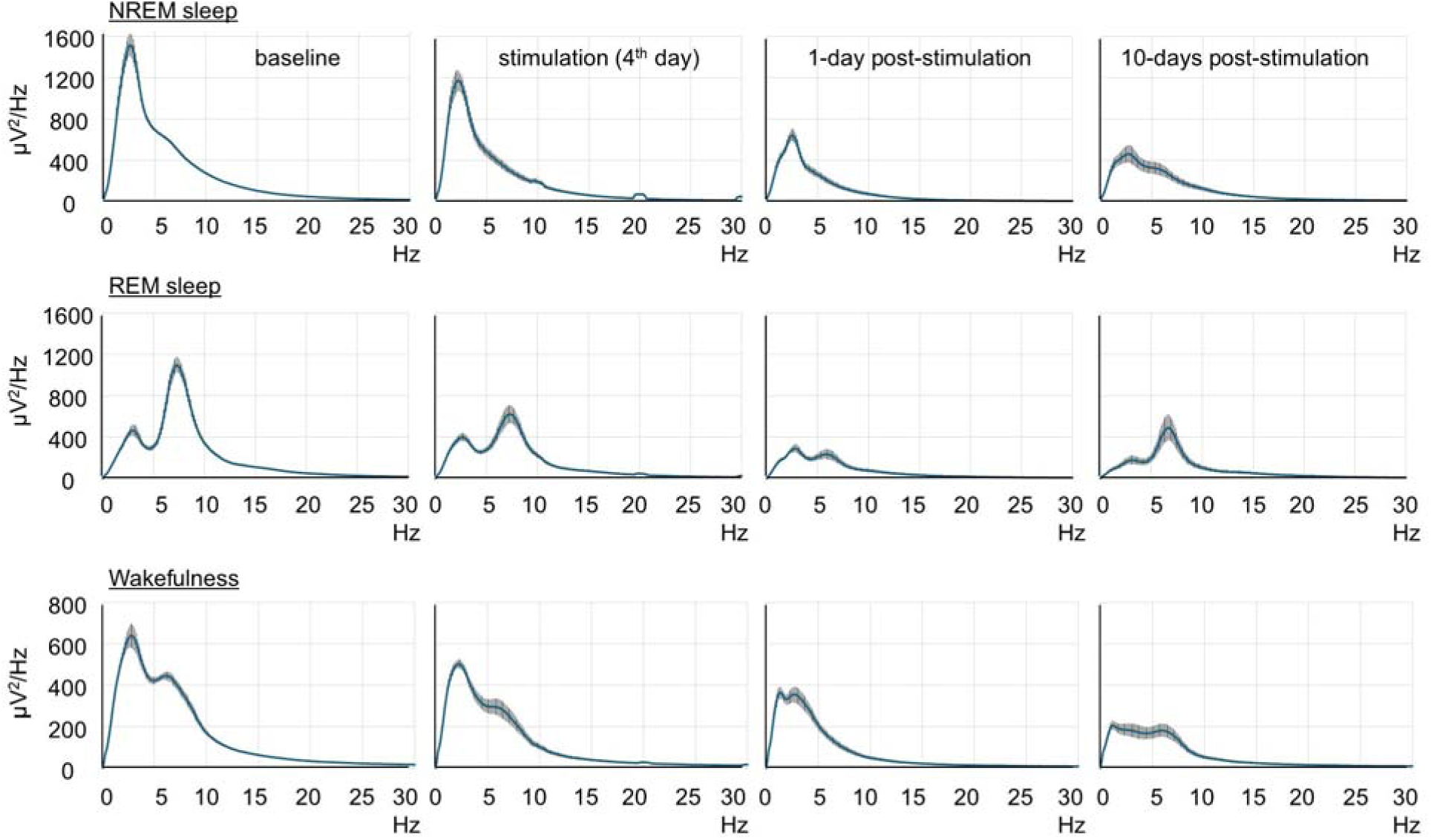
Effect of brain exposure to red light on sleep amounts and EEG power in 16-month-old mice. EEG spectral power (blue lines) during NREM sleep (Top), REM sleep (Middle) and wakefulness (Bottom) in mice calculated during baseline, stimulation and post-stimulation days. Error bars indicate the standard error of the mean.

**Extended Data Fig. 4.**
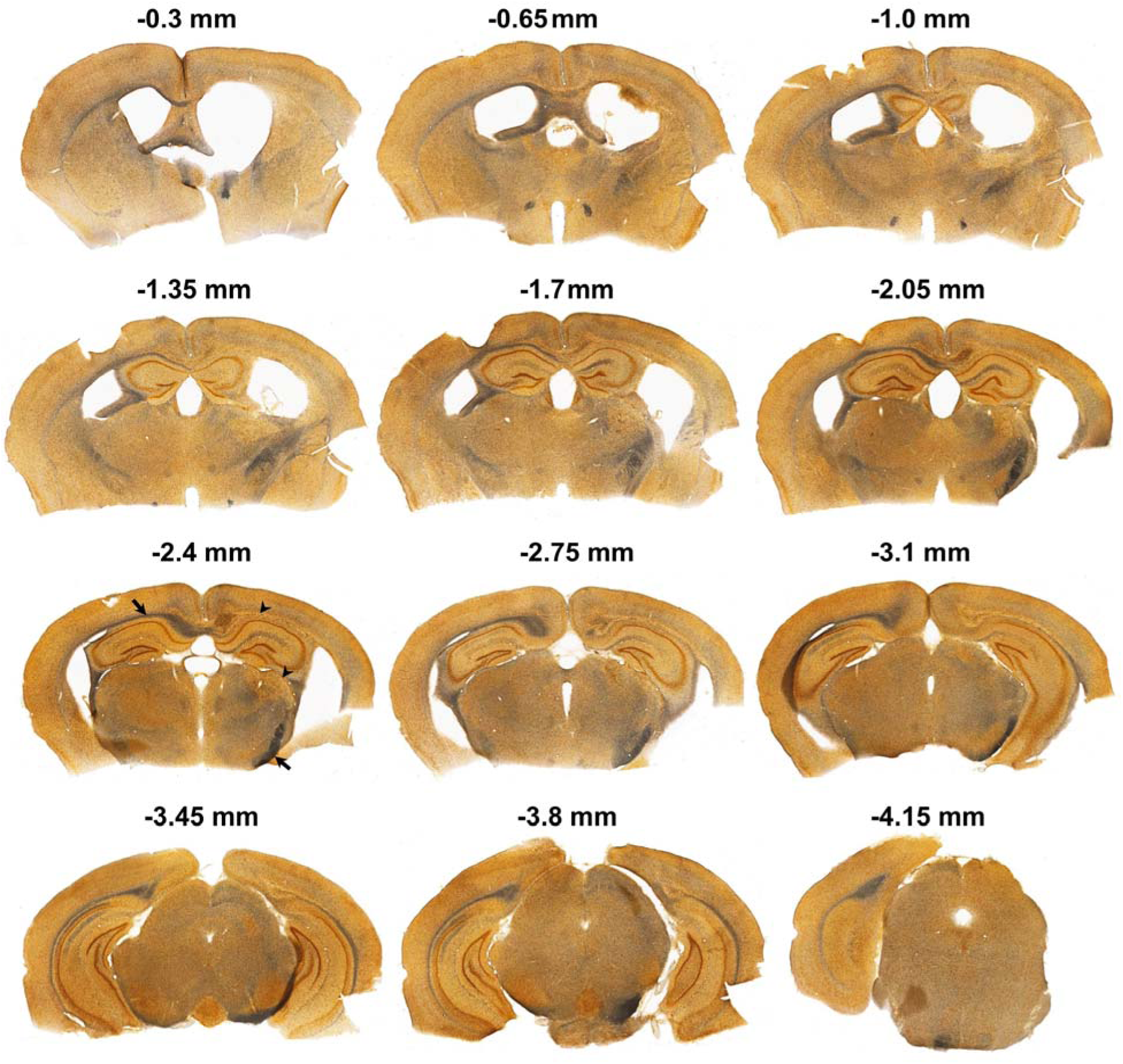
Axonal damage following brain exposure to red light assessed with silver staining in a 16-month-old mouse. Representative images of silver-stained coronal sections are shown at several locations relative to bregma. LED was located above the cerebral cortex at the right hemisphere about 2-2.5 mm posterior to bregma. White matter areas with prominent silver staining are indicated by black arrows. A damaged areas with absent silver staining are indicated by black arrowheads.

**Extended Data Fig. 5.**
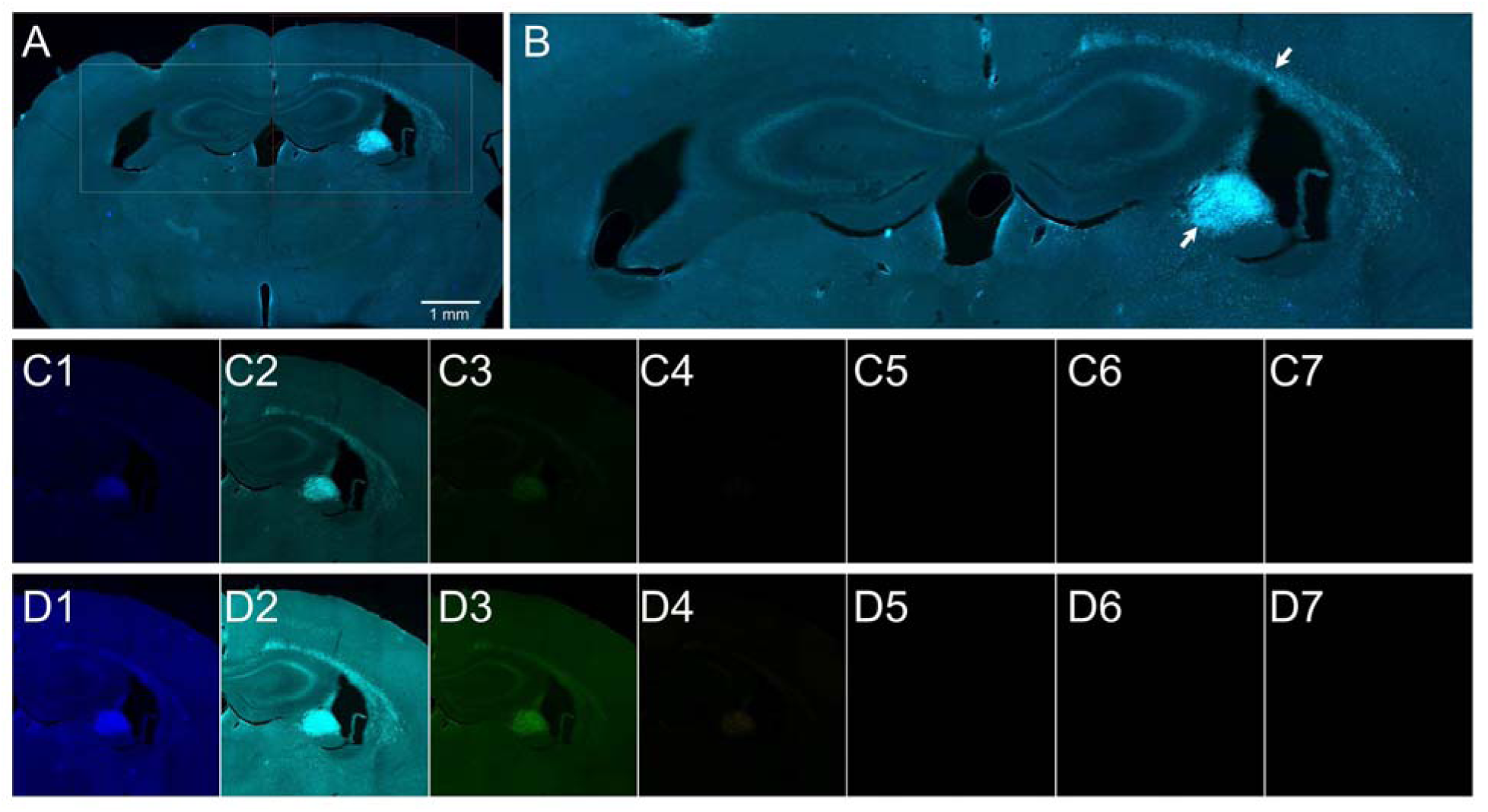
Autofluorescence in the brain tissue of a 16-month-old mouse assessed by multispectral imaging. (**A**) Brain section of a 16-month-old mouse stimulated with red light. (**B**) The area outlined in panel A at a higher modification. Autofluorescent areas are indicated by white arrows. (**C1**) Opal DAPI imaging filter, excitation 368 nm, emission 461nm, (**C2**) Opal 480 imaging filter, excitation 450 nm, emission 500 nm, (**C3**) Opal 520 imaging filter, excitation 494 nm, emission 525 nm, (**C4**) Opal 570 imaging filter, excitation 550 nm, emission 570 nm, (**C5**) Opal 620 imaging filter, excitation 588 nm, emission 616 nm, (**C6**) Opal 690 imaging filter, excitation 676 nm, emission 694 nm, (**C7**) Opal 780 imaging filter, excitation 750 nm, emission 770 nm, (**D1** – **D7**) The same imaging filters as in C1 – C7, but the brightness increased 150%.

**Extended Data Fig. 6.**
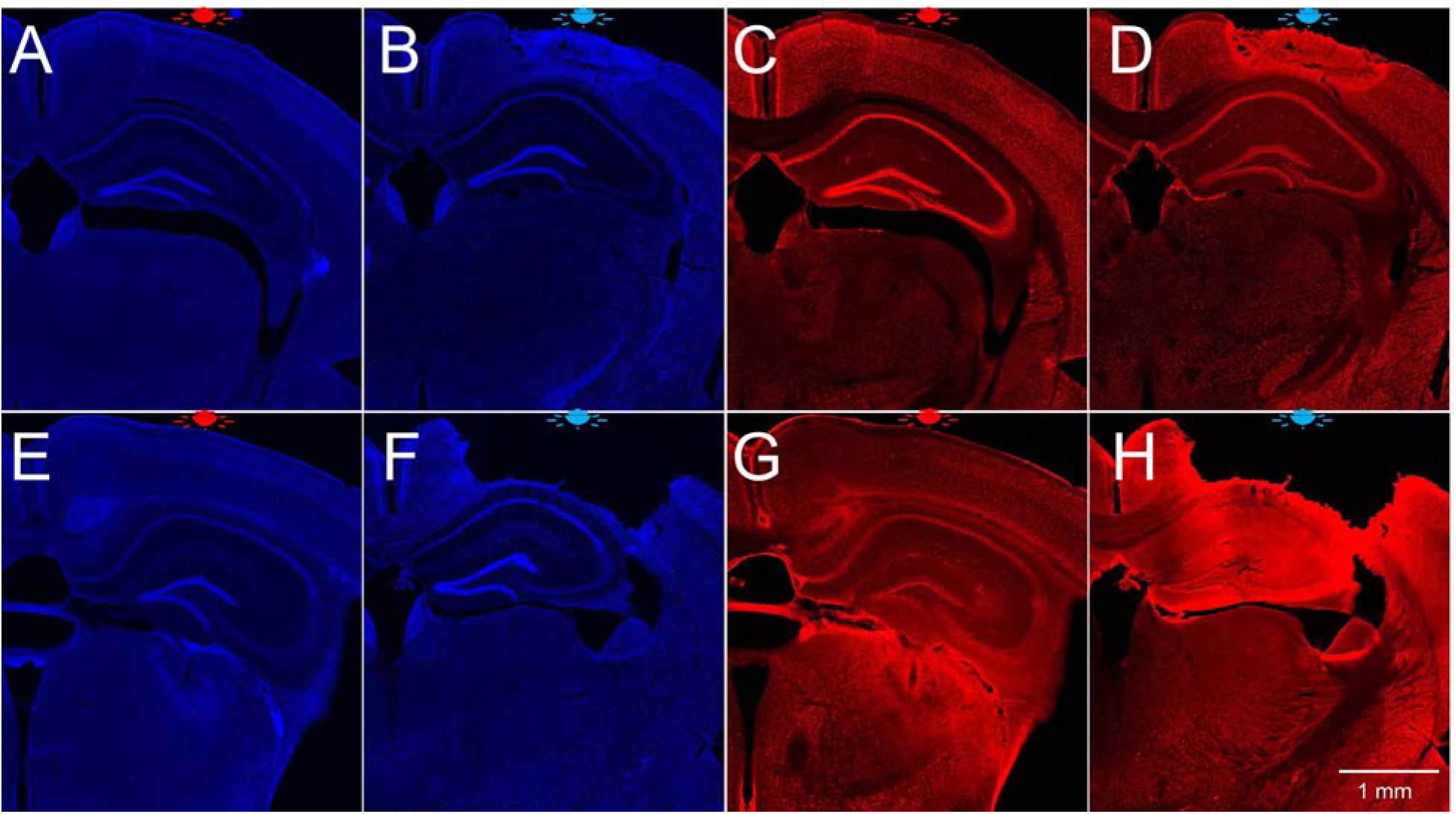
DAPI staining and NeuN immunostaining in the brain of young and old mice exposed to red and blue light. (**A**) DAPI staining of the brain of 3-month-old mouse exposed to red light, (**B**) DAPI staining of the brain of 3-month-old mouse exposed to blue light, (**C**) NeuN immunostaining of the brain of 3-month-old mouse exposed to red light, (**D**) NeuN immunostaining of the brain of 3-month-old mouse exposed to blue light, (**E**) DAPI staining of the brain of 16.5-month-old mouse exposed to red light, (**F**) DAPI staining of the brain of 14-month-old mouse exposed to blue light, (**G**) NeuN immunostaining of the brain of 16.5-month-old mouse exposed to red light, (**H**) NeuN immunostaining of the brain of 14-month-old mouse exposed to blue light. Approximate location of LED is shown at the top of each panel.

**Extended Data Fig. 7.**
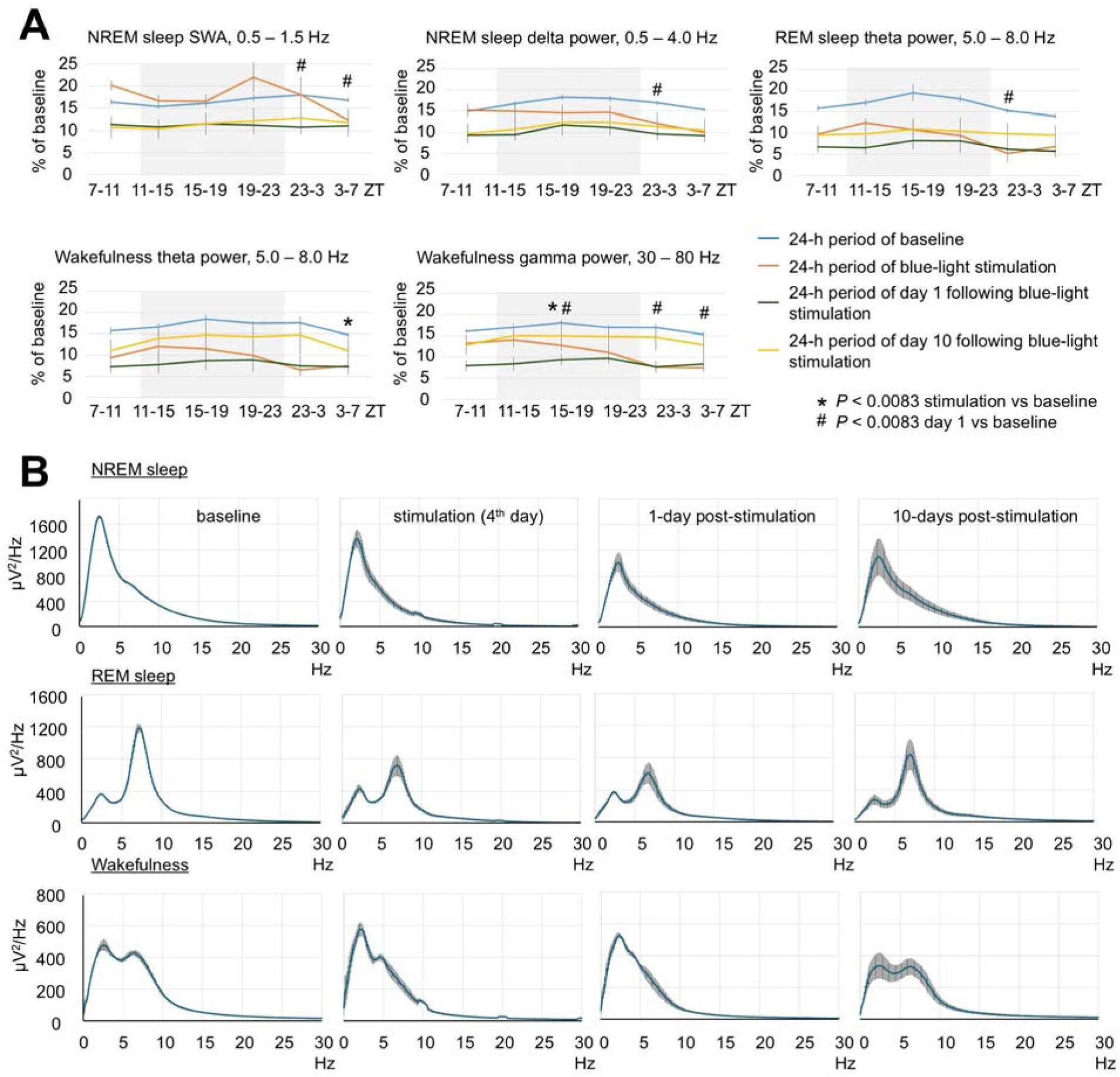
Effect of brain exposure to blue light on sleep amounts and EEG power in 14-month-old mice. (**A**) Profile of changes of EEG power in SWA, delta, theta and gamma range in mice exposed to blue light. Each datapoint represents power calculated during a 4-hour period. One-way repeated measures ANOVA followed by Tukey’s post hoc test, *#†*P* < 0.0083 vs baseline. (**B**) Spectral distribution of EEG power density in the frequencies between 0.5 and 30 Hz during NREM sleep (Top), REM sleep (Middle) and wakefulness (Bottom) in mice calculated during baseline, stimulation and post-stimulation days. Error bars indicate the standard error of the mean.

**Extended Data Fig. 8.**
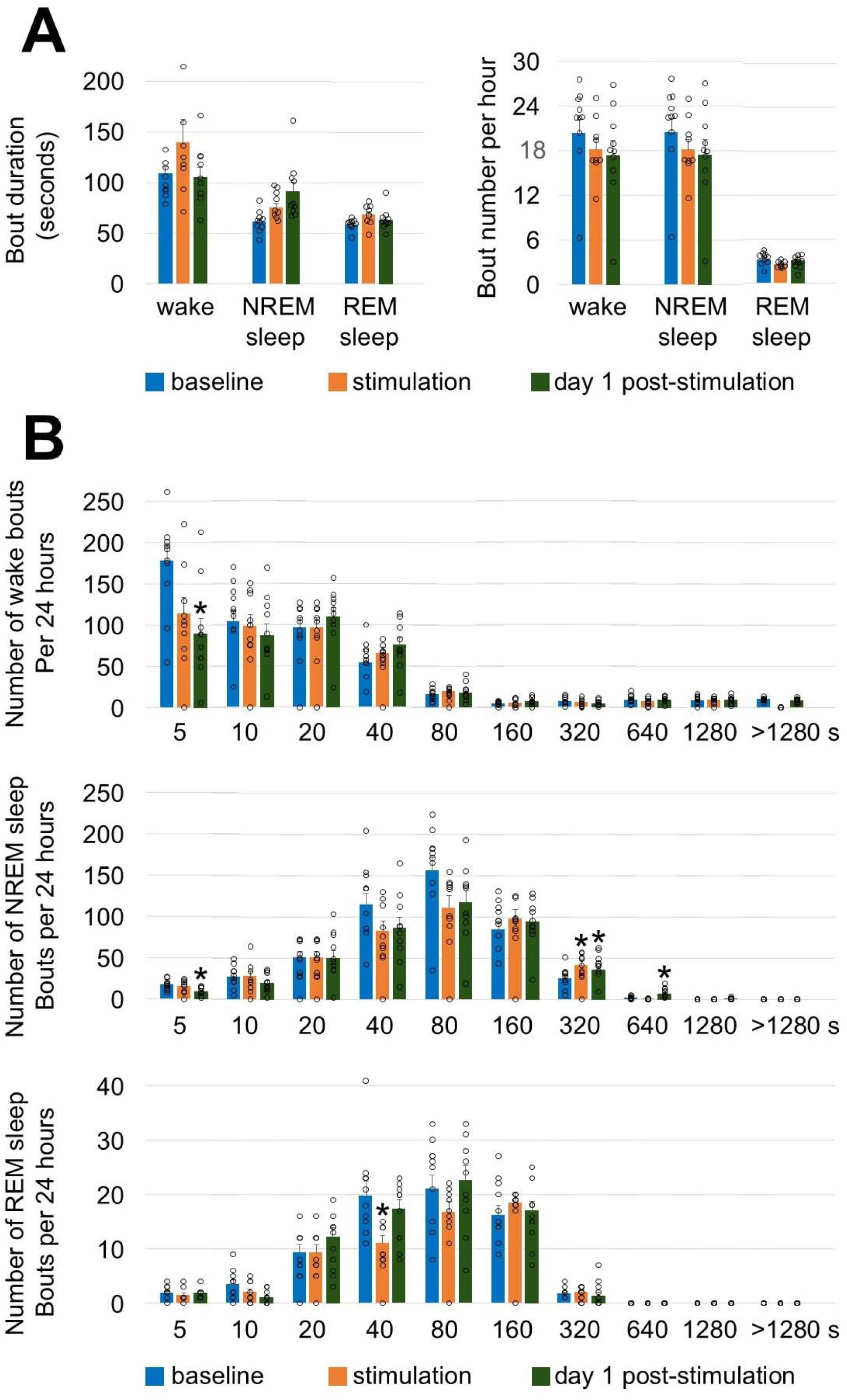
Effect of brain exposure to red light on sleep and wakefulness architecture in 7.5-month-old mice. (**A**) Average bout duration and bout number calculated during 24-h period of baseline, stimulation, and post-stimulation days. (**B**) Episode frequency histograms of wakefulness (upper panel), NREM (middle panel) and REM sleep (lower panel) during baseline, stimulation, and post-stimulation days. Episodes were partitioned in ten exponentially increased duration bins (upper bin limit from 5s to > 1280s) Error bars indicate the standard error of the mean. One-way repeated measures ANOVA followed by Tukey’s post hoc test. \**P*<0.05 vs baseline.

**Extended Data Fig. 9.**
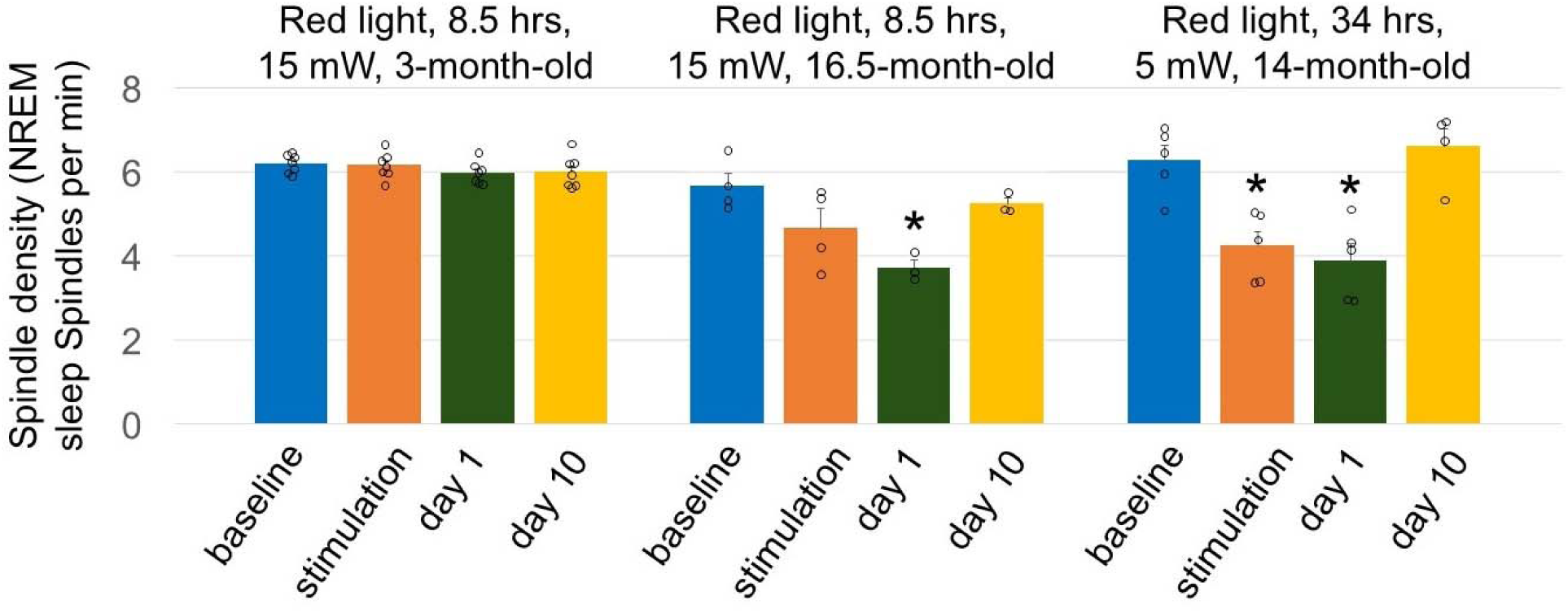
Effect of brain exposure to red light on sleep spindle density. Mean spindle density calculated during 24 hrs of baseline, stimulation and post-stimulation day 1 and 10 in young and old mice. Error bars indicate the standard error of the mean. One-way repeated measures ANOVA followed by Tukey’s post hoc test. \**P*< 0.05 vs baseline.

**Extended Data Fig. 10.**
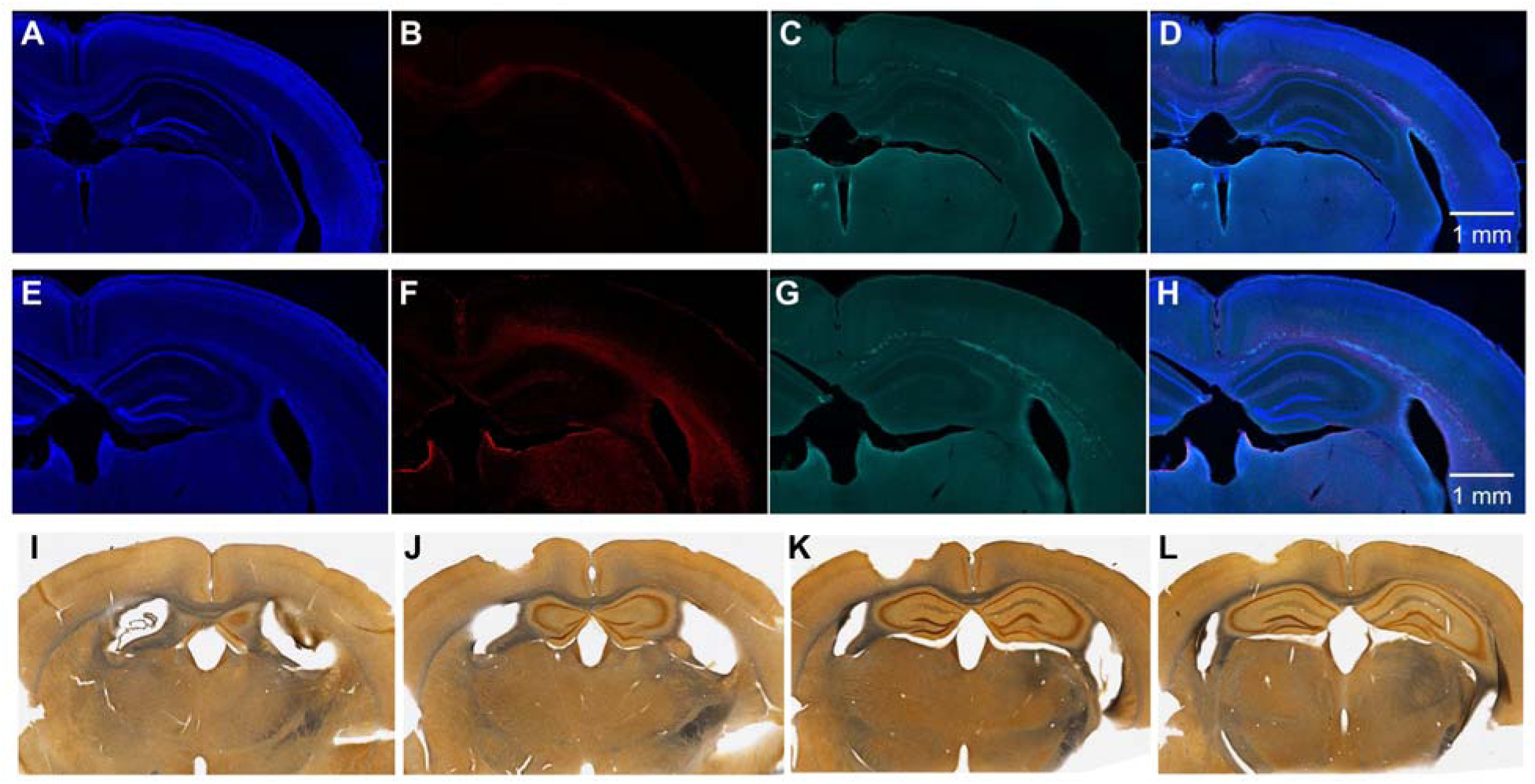
Damage in the brain of a 15-month-old mouse exposed to red light delivered at lower power and longer duration. (**A**) DAPI staining, (**B**) GFAP staining, (**C**) autofluorescence under green filter, and (**D**) merged image of A-C in a mouse exposed to red light. (**E**) DAPI staining, (**F**) IBA-1 staining, (**G**) autofluorescence under green filter, and (**H**) merged image of E-G in a mouse exposed to blue light. Silver staining in brain sections located approximately at (**I**) -0.7 mm, (**J**) -1.3 mm, (**K**) -1.9 mm, (**L**) -2.5 mm from bregma.

## References

1. Emiliani, V. et al. Optogenetics for light control of biological systems. Nat. Rev. Methods Primers. 2, (2022).

2. Lehtinen, K., Nokia, M.S. & Takala, H. Red Light Optogenetics in Neuroscience. Front Cell Neurosci. 15, 778900 (2021).

3. Tyssowski, K.M. & Gray, J.M. Blue Light Increases Neuronal Activity-Regulated Gene Expression in the Absence of Optogenetic Proteins. eNeuro. 6, (2019).

4. Al Juboori, S.I. et al. Light scattering properties vary across different regions of the adult mouse brain. PLoS. ONE. 8, e67626 (2013).

5. da Rocha, R.B. et al. The role of light emitting diode in wound healing: A systematic review of experimental studies. Cell Biochem. Funct. 42, e4086 (2024).

6. Wells, A., Rigby, J., Castel, C. & Castel, D. Pulsed Red and Blue Photobiomodulation for the Treatment of Thigh Contusions and Soft Tissue Injury: A Randomized Controlled Trial. J Sport Rehabil. 33, 20–26 (2024).

7. DuBois, D.W. et al. Maintenance of optogenetic channel rhodopsin (ChR2) function in aging mice: Implications for pharmacological studies of inhibitory synaptic transmission, quantal content, and calcium homeostasis. Neuropharmacology 238, 109651 (2023).

8. Chong, H.R., Ranjbar-Slamloo, Y., Ho, M.Z.H., Ouyang, X. & Kamigaki, T. Functional alterations of the prefrontal circuit underlying cognitive aging in mice. Nat. Commun. 14, 7254 (2023).

9. Bang, E., Fincher, A.S., Nader, S., Murchison, D.A. & Griffith, W.H. Late-Onset, Short-Term Intermittent Fasting Reverses Age-Related Changes in Calcium Buffering and Inhibitory Synaptic Transmission in Mouse Basal Forebrain Neurons. J Neurosci. 42, 1020–1034 (2022).

10. Sun, X.Y. et al. Two parallel medial prefrontal cortex-amygdala pathways mediate memory deficits via glutamatergic projection in surgery mice. Cell Rep. 42, 112719 (2023).

11. Yang, J. et al. Chronic blue light leads to accelerated aging in Drosophila by impairing energy metabolism and neurotransmitter levels. Front Aging 3, 983373 (2022).

12. Thankachan, S., Gerashchenko, A., Kastanenka, K.V., Bacskai, B.J. & Gerashchenko, D. Optimization of real-time analysis of sleep-wake cycle in mice. MethodsX. 9, 101811 (2022).

13. Nairuz, T., Sangwoo, C. & Lee, J.H. Photobiomodulation Therapy on Brain: Pioneering an Innovative Approach to Revolutionize Cognitive Dynamics. Cells 13, (2024).

14. Salehpour, F. et al. Brain Photobiomodulation Therapy: a Narrative Review. Mol. Neurobiol. 55, 6601–6636 (2018).

15. Avvenuti, G. et al. Integrity of Corpus Callosum Is Essential for theCross-Hemispheric Propagation of Sleep Slow Waves:A High-Density EEG Study in Split-Brain Patients. J Neurosci. 40, 5589–5603 (2020).

16. Gudberg, C. et al. Individual differences in slow wave sleep architecture relate to variation in white matter microstructure across adulthood. Front Aging Neurosci. 14, 745014 (2022).

17. Buzsaki, G., Gage, F.H., Czopf, J. & Bjorklund, A. Restoration of rhythmic slow activity (theta) in the subcortically denervated hippocampus by fetal CNS transplants. Brain Res. 400, 334–347 (1987).

18. Blasi, F., Whalen, M.J. & Ayata, C. Lasting pure-motor deficits after focal posterior internal capsule white-matter infarcts in rats. J Cereb. Blood Flow Metab 35, 977–984 (2015).

19. Suhan, S. et al. Experimental assessment of the safety and potential efficacy of high irradiance photostimulation of brain tissues. Sci. Rep. 7, 43997 (2017).

20. Chen, R. et al. Deep brain optogenetics without intracranial surgery. Nat. Biotechnol. 39, 161–164 (2021).

21. Stujenske, J.M., Spellman, T. & Gordon, J.A. Modeling the Spatiotemporal Dynamics of Light and Heat Propagation for In Vivo Optogenetics. Cell Rep. 12, 525-534 (2015).

22. Kasprzycka, W., Szumigraj, W., Wachulak, P. & Trafny, E.A. New approaches for low phototoxicity imaging of living cells and tissues. Bioessays 46, e2300122 (2024).

23. Francon, A. et al. Phototoxicity of low doses of light and influence of the spectral composition on human RPE cells. Sci. Rep. 14, 6839 (2024).

24. Zein, R., Selting, W. & Hamblin, M.R. Review of light parameters and photobiomodulation efficacy: dive into complexity. J Biomed. Opt. 23, 1–17 (2018).

25. Fernandes, F. et al. Devices used for photobiomodulation of the brain-a comprehensive and systematic review. J Neuroeng. Rehabil. 21, 53 (2024).

26. Carneiro, A.M.C. et al. Transcranial Photobiomodulation Therapy in the Cognitive Rehabilitation of Patients with Cranioencephalic Trauma. Photobiomodul. Photomed. Laser Surg. 37, 657–666 (2019).

27. Hipskind, S.G. et al. Pulsed Transcranial Red/Near-Infrared Light Therapy Using Light-Emitting Diodes Improves Cerebral Blood Flow and Cognitive Function in Veterans with Chronic Traumatic Brain Injury: A Case Series. Photobiomodul. Photomed. Laser Surg. 37, 77–84 (2019).

28. Ma, L.Z. et al. Time spent in outdoor light is associated with the risk of dementia: a prospective cohort study of 362094 participants. BMC. Med 20, 132 (2022).

29. Li, H., Cui, F., Wang, T., Wang, W. & Zhang, D. The impact of sunlight exposure on brain structural markers in the UK Biobank. Sci. Rep. 14, 10313 (2024).

30. Song, Y. et al. Age-dependent effects of blue light exposure on lifespan, neurodegeneration, and mitochondria physiology in Drosophila melanogaster. NPJ. Aging 8, 11 (2022).

31. Somasundaram, I. et al. Mitochondrial dysfunction and its association with age-related disorders. Front Physiol 15, 1384966 (2024).

32. Li, W. et al. Boosting neuronal activity-driven mitochondrial DNA transcription improves cognition in aged mice. Science 386, eadp6547 (2024).

33. Lin, M.T. & Beal, M.F. Mitochondrial dysfunction and oxidative stress in neurodegenerative diseases. Nature 443, 787–795 (2006).

34. Yabluchanskiy, A. et al. Age-related alterations in the cerebrovasculature affect neurovascular coupling and BOLD fMRI responses: Insights from animal models of aging. Psychophysiology 58, e13718 (2021).

35. Ilie, O.D. et al. Mini-Review on Lipofuscin and Aging: Focusing on The Molecular Interface, The Biological Recycling Mechanism, Oxidative Stress, and The Gut-Brain Axis Functionality. Medicina (Kaunas.) 56, (2020).

## References

36. Leopold, A.V., Thankachan, S., Yang, C., Gerashchenko, D. & Verkhusha, V.V. A general approach for engineering RTKs optically controlled with far-red light. Nat. Methods 19, 871–880 (2022).

37. Prerau, M.J., Brown, R.E., Bianchi, M.T., Ellenbogen, J.M. & Purdon, P.L. Sleep Neurophysiological Dynamics Through the Lens of Multitaper Spectral Analysis. Physiology. (Bethesda.) 32, 60–92 (2017).

38. Babadi, B. & Brown, E.N. A review of multitaper spectral analysis. IEEE Trans. Biomed. Eng 61, 1555–1564 (2014).

39. Uygun, D.S. et al. Validation of an automated sleep spindle detection method for mouse electroencephalography. Sleep 42, (2019).

40. Thankachan, S. et al. Thalamic Reticular Nucleus Parvalbumin Neurons Regulate Sleep Spindles and Electrophysiological Aspects of Schizophrenia in Mice. Sci. Rep. 9, 3607 (2019).

